# Inducible formation of leading cells driven by CD44 switching gives rise to collective invasion

**DOI:** 10.1101/387092

**Authors:** Cuixia Yang, Manlin Cao, Yiwen Liu, Yiqing He, Yan Du, Guoliang Zhang, Feng Gao

## Abstract

Collective invasion into adjacent tissue is a hallmark of luminal breast cancer, with about 20% of cases that eventually undergo metastasis. It remained unclear how less aggressive luminal-like breast cancer transit to invasive cancer. Our study revealed that CD44^hi^ cancer cells are the leading subpopulation in collective invading cancer cells, which could efficiently lead the collective invasion of CD44^lo^/follower cells. CD44^hi^/leading subpopulation showed specific gene signature of a cohort of hybrid epithelial/mesenchymal state genes and key functional co-regulators of collective invasion, which was distinct from CD44^lo^/follower cells. However, the CD44^hi^/leading cells, in partial-EMT state, were readily switching to CD44^lo^ phenotype along with collective movements and vice versa, which is spontaneous and sensitive to tumor microenvironment. The CD44^lo^-to-CD44^hi^ conversion is accompanied with a shift of CD44s-to-CD44v, but not corresponding to the conversion of non-CSC-to-CSC. Therefore, the CD44^hi^ leader cells are not a stable subpopulation in breast tumors. This plasticity and ability to generate CD44^hi^ carcinoma cells with enhanced invasion-initiating powers might be responsible for the transition from in situ to invasive behavior of luminal-type breast cancer.

**Significance:** Now, the mechanisms involved in local invasion and distant metastasis are still unclear. We identified a switch of CD44 that drives leader cell formation during collective invasion in luminal breast cancer. We provided evidence that interconversions between low and high CD44 states occur frequently during collective invasion. Furthermore, these findings demonstrated that the CD44^hi^/leader cells featuring partial EMT are inducible and attainable in response to tumor microenvironment. The CD44^lo^ cancer cells are plastic that readily shift to CD44^hi^ state, accompanied with shifts of CD44s-to-CD44v, thereby increasing tumorigenic and malignant potential. There are many “non-invasiveness” epithelial/follower cells with reversible invasive potential within an individual tumor, that casting some challenges on molecular targeting therapy.

## Introduction

The challenge to understand the nature of tumor cells that initiate metastases is of great clinical significance. Invasion into adjacent tissue often followed by metastasis is a hallmark of breast cancer (BrCa). Growing evidence showed that the metastatic dissemination of epithelial breast tumors can occur in a manner as collective cells (Friedl et al, 2012) without activation of the epithelial-mesenchymal transition (EMT) program(Fischer et al, 2015; Zheng et al, 2015), in which cancer cells locally invade the peritumoral stroma as cohesive clusters while maintaining cell-cell junctions, even traveling as clustering circulating-tumor cells (Aceto et al, 2014; Yu et al, 2013)and more efficiently seeding to form metastases than single circulating tumor cells(Cheung et al, 2016; Zajac et al, 2018). EMT models could not adequately describe the invasive behaviors of breast cancer cells in collective movement(Shamir et al, 2014). The nature of the cancer cells that give rise to cohesive invading is still poorly understood.

At the onset of collective cell invasion, a subset of cancer cells appear at the leading edge acquires a distinct phenotype with characteristic morphology and motility. It was revealed that basal genes, such as cytokeratin 14 (K14) or p63(Cheung et al, 2013), and some signaling moleculars, such as Erk1/2 and RhoA located at the leading front cells(Chapnick & Liu, 2014; Reffay et al, 2014). Those leading cells can facilitate invasion by forming invasive protrusions, which provide traction and trigger modifications of the ECM in leader-follower cooperative manner(Nguyen-Ngoc et al, 2012). It is more of challenge for epithelial cancers to drive leader cell formation and initiate cell spread in a collective manner. The ability to acquire a distinct “leader” phenotype varies among different tumor cell populations originally derived from different patients with individual intracellular and extracellular microenvironment (Liu et al, 2013)’ (Dang et al, 2011; Nguyen-Ngoc et al, 2012). The mechanisms of how collectively cell invading into the surroundings was triggered are still not clearly understood.

Clinically, breast cancers are heterogeneous, often containing multiple subpopulations of cancer cells with distinct molecular states(Almendro et al, 2013; McGranahan & Swanton, 2015), which are likely to be plastic and adaptive to microenvironment stimuli. The heterogeneity and plasticity in mammary epithelial cell may cause poor response to endocrine therapy (Ito et al, 2015) and high risk of metastasis(Kennecke et al, 2010), although a more favorable prognosis could be expected in most of luminal breast cancers. Understanding how these subpopulations in luminal epithelial cells, though displaying less invasive than basal like BrCa cells, acquire phenotypic changes and enable an enhanced invasive ability to metastasis is of great importance.

Here, we investigated the switch of phenotypic states in subpopulations in epithelial breast cancer induced by the shift of CD44 levels during collective cell invasion. The phenotypes of cancer cell were showed to be dynamic and plastic(Friedl & Alexander, 2011; Matak et al, 2017), and a dynamic conversion between CSCs and Non-CSCs was also revealed(Iliopoulos et al, 2011). CD44, as a mesenchymal marker, was used to identify cancer stem cells, and also contributed to regulate tumor stemness and metastasis(Ahmed et al, 2016). Interconversions between CSCs and Non-CSCs were usually happened that illustrating the dynamic and plastic of cancer cell phenotypes(Friedl & Alexander, 2011; Matak et al, 2017)^,23^. Switch in expression pattern of CD44 standard (CD44s) and variant isoforms (CD44v) has been shown to influence EMT process, but functional significances of CD44 variants are not yet fully understood(Brown et al, 2011; Preca et al, 2015). Nevertheless, the involvement of CD44 in leader cell formation during epithelial collective invasion has not been studied. We therefore observed the inducible and exchangeable acquirement of CD44 along with other leader cell features, and monitored the formation of leading tips during collective migration. Our results revealed that the plasticity and ability of epithelial cancer cells, CD44^lo^-to-CD44^hi^ conversion, generate invasive leading cells, which might be contribute to the transition from in situ to invasive behavior of luminal-type BrCas.

## Results

### 1. CD44 is a featured molecular of leading cells during collective invasion in breast cancers

To investigate intratumor heterogeneity related to CD44 expression during progression in breast carcinoma, we performed immunohistochemistry analysis of 247 human breast tumors of different molecular subtypes, 30 of which containing the tumor-stromal interface of the invasive edge. We analyzed the tumors for the expression of CD44 in 41 luminal A, 92 luminal B, 44 HER2+, and 70 triple negative (TN), as defined by immunohistochemical analysis. Interestingly, at the tumor-stromal interface in all four molecular types, multicellular groups of cells were revealed invading into the adjacent stroma (Fig. 1A). Remarkably, the level of CD44 was significantly upregulated in the edge of leading cells in the invading strands, but not in other cells in the inner region of the collective clusters (Fig. 1A). The frequency of CD44^hi^ cancer cells is correlated with the clinical molecular subtypes, displaying the highest in triple negative and the lowest in luminal-like cases examined (Fig. 1B and Supplemental Table S1), which is consistent with previous studies(Park et al, 2010). No significant differences were detected when the level of CD44^hi^ cell was compared between luminal-A and luminal-B cases (P = 0.9301) (Fig. 1B and Supplemental Table S1).

**Fig.1.**
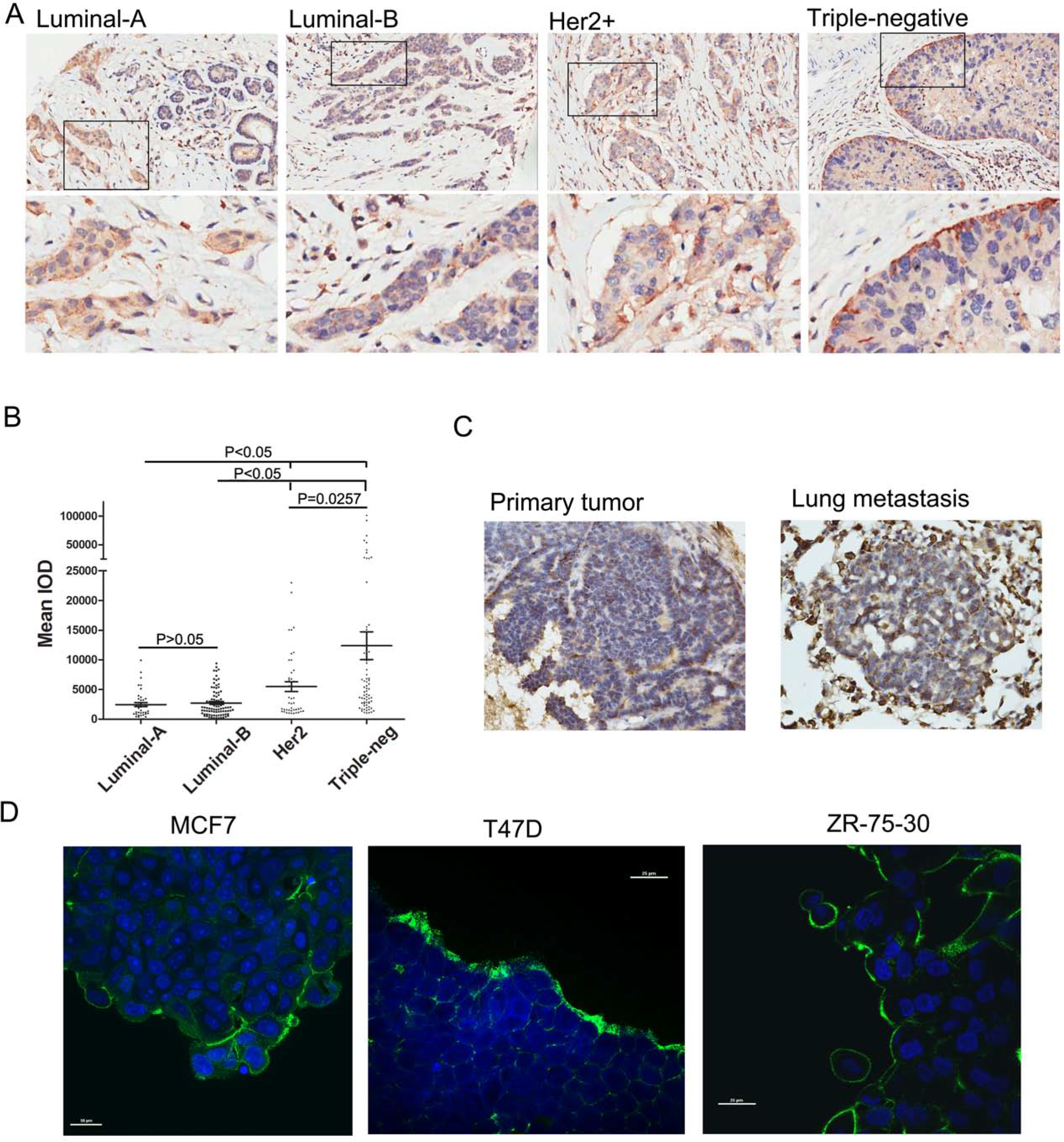
CD44hi cancer cells are enriched at the leading edge in collective invasive tumor cells and in lung metastases. (A) Immunohistochemical expression of CD44 in mammary tumor sections based on molecular subtypes. CD44 with homogenous membranous staining is enriched in the leading breast carcinoma cells. Arrow, leading CD44hi cancer cell with forked protrusions. (B) The expression of CD44 in different molecular subtypes of breast cancers. (C) CD44hi cancer cells are enriched at the invasive border in primary tumor and in lung metastases from MMTV-PyMT mice. (D)Expression of CD44 in luminal breast cancer cells during collective migration. Scale bars, 25μm.

Similar to clinical tumor tissues, in orthotopic xenografts in mice, CD44 was also enriched at the invading boundary in luminal-like primary breast tumor and lung metastases (Fig. 1C). Also, immunostaining assay further indicated that cancer cells with CD44 high levels were located at migrating tips during collective movements in luminal-like breast cancer cells (Fig. 1D and Fig.S1). Follower cells at the inner region during collective migration, however, presented only low levels of CD44. Taken together, the distinctively high levels of CD44 in the invading front observed indicate that the CD44^hi^ state is one of molecular features of leader cells, and breast tumors are composed of subpopulations of cancer cells with distinct CD44 expressions.

### 2. Oncogenic properties of CD44^lo^ and CD44^hi^ cancer cells generated during the collective invasion in vivo

Clinically, 20% of luminal breast cancers finally metastasis to distant organs^20^. To test the role of CD44^hi^ status on the leading edge cells in luminal breast cancers, though with general lower-level of CD44 than basal cancers (Fig. 1B and Table S1), we next sought to determine the function of CD44^lo^ and CD44^hi^ cells arising in vivo in collective invasion. Purified subpopulations of CD44^hi^ and CD44^lo^ cells from luminal-like tumors were isolated by fluorescence activated cell sorting (FACS). A subsequent cell proliferation assay displayed that CD44^hi^ cells are significantly more proliferative than CD44^lo^ subsets (Fig. 2A). Compatible with this result, an EdU incorporation assay also revealed that the proliferation of CD44^hi^ cells was higher than that of CD44^lo^ cells (Fig. 2B). Furthermore, we investigated the invasiveness of each of the subpopulations using the Matrigel-coated transwell filter assay, and the CD44^hi^ populations showed by far more invasive (Fig. 2C). We also determined stem/progenitor activity by mammosphere formation assay. Results showed that CD44^hi^ subpopulation is more capable of forming mammospheres compared with CD44^lo^ subsets (Fig. 3D). Interestingly, the CD44^lo^ population is able to form mammospheres too, although with less efficient.

**Fig.2.**
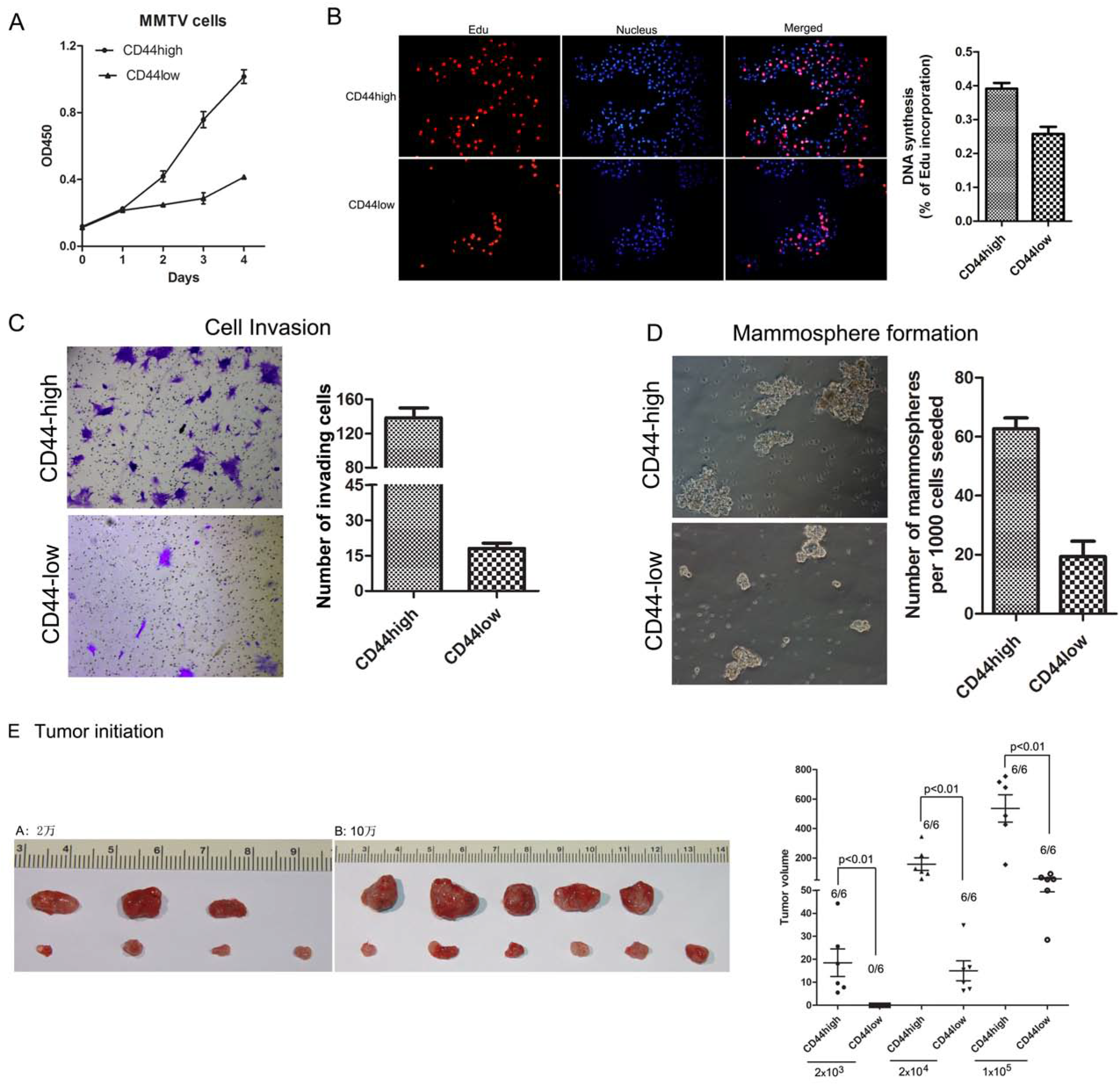
The CD44hi and CD44lo cells generated during the collective invasion have distinct functions. (A) Proliferation of FACS-sorted EpCAM^+^/CD45^-^/CD44^hi^ or EpCAM^+^/CD45^-^/CD44^lo^° cancer cells from MMTV/PyMT mice assessed using a CCK-8 kit at the indicated days. Data are the means±s.e.m. of three experiments.(B) DNA synthesis assessed using an EdU (5-ethynyl-2’-deoxyuridine) assay in sorted EpCAM^+^/CD45^-^/CD44^hi^ and EpCAM^+^/CD45^-^/CD44^lo^° cancer cells. Cells were fluorescently stained with EdU (red). Nuclei were stained with DAPI (blue). Micrographs represent at least three experiments. Scale bar, 200 mm. Quantitative EdU assay data are the means±s.e.m. of three experiments. *p < 0.05, **p < 0.01 (Student’s t-test).(C)Invasion assay were performed using sorted EpCAM^+^/CD45^-^/CD44^hi^ or EpCAM^+^/CD45^-^/CD44^lo^ cells and allowed to migrate for 24 h. Representative images are shown, scale bar=200 μm(n=2 independent cell cultures and experiments, 3 technical replicates of each). Throughout figure, all graphs are shown as mean±s.e.m. *p < 0.05; **p < 0.01 by two-tailed Student’s t-test. (D) Mammosphere-forming capacity of CD44^hi^ and CD44^lo^ populations. CD44hi or CD44lo cells (1 × 10^3^) were sorted into each well of 24-well ultra-low attachment dishes containing mammosphere growth medium. Cells were cultured for 12 days, and the photos were taken using a light microscope. Scale bars: 100 μm. Numbers of mammospheres per well were quantified. *p < 0.01 compared with CD44^lo^ The experiment was performed in triplicate. (E) Assessment of the tumorigenicity of the EpCAM+/CD45-/CD44^hi^ or EpCAM^+^/CD45^-^/CD44^lo^° cells (sorted from MMTV/PyMT mice) orthotropically injected into NOD-SCID mice at several dilutions with 200-5×10^5^ cell, as determined by tumor volume, is shown. Numbers indicate the frequency of tumor formation. Dot plots indicate individual tumor volumes (Mean indicated by solid line). *p < 0.05 compared with CD44^lo^ cells-derived tumors.

**Fig.3.**
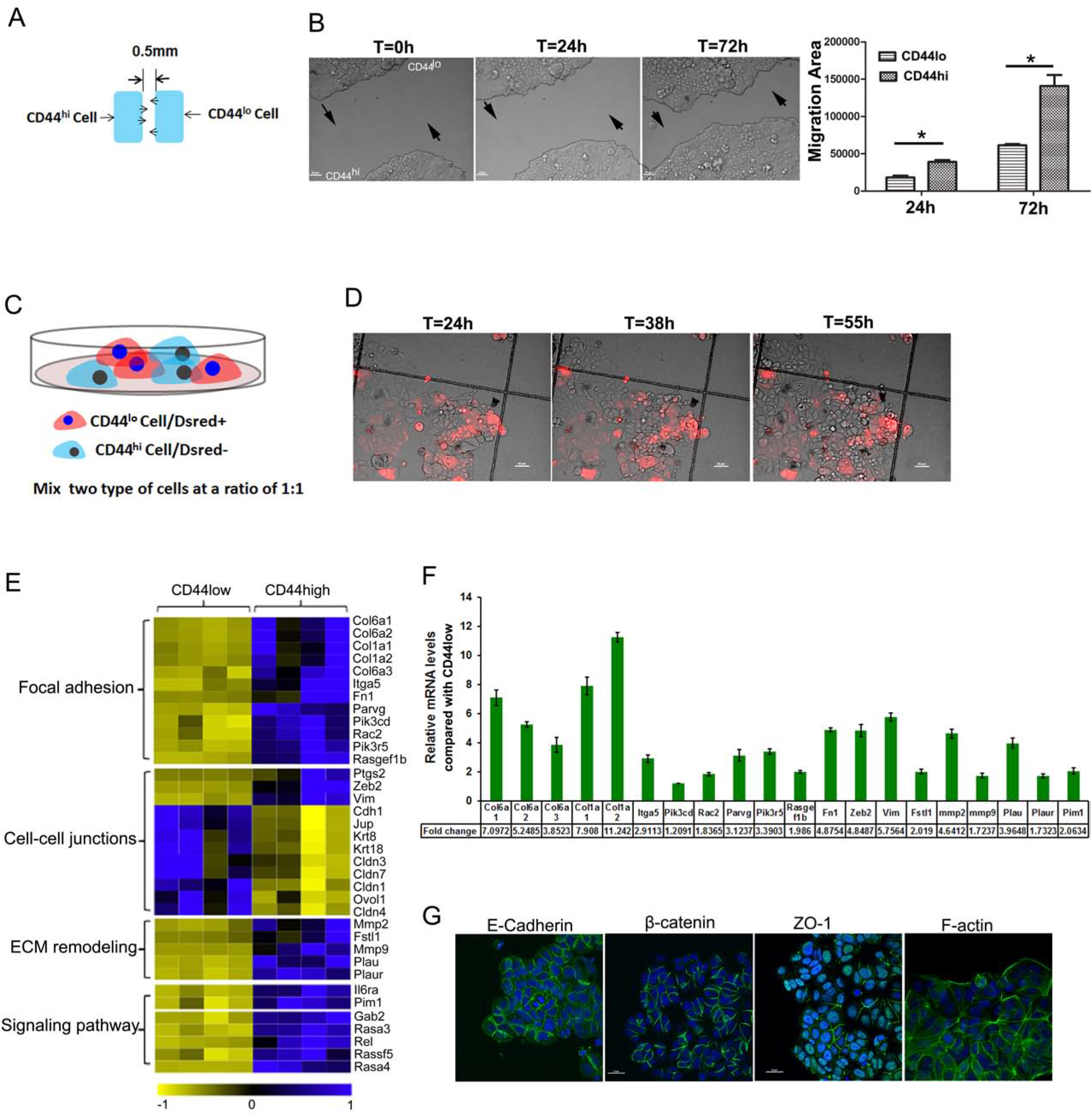
CD44^hi^ cancer cells with the invasive phenotype lead the luminal CD44^lo^ cancer cells during the collective invasion. (A) Schematic representation of the in vitro collective migrating model in an epithelial monolayer. The FACS-sorted CD44^hi^ cells and CD44^lo^ cancer cells were cultured within two confined wells separately. After confluency, the confinement was removed to trigger epithelial cluster migration. The space between two cell monolayers was 500μm wide. (B)Phase contrast microscopy images of the forward facing migration of CD44^hi^ cells and CD44^lo^ epithelial monolayers. Scale bars, 50μm. (C) Schematic representation of the collective migration with the mixed two types of cells. (D) CD44^hi^/Dsred+ cells migrate collectively and lead trailing CD44^lo^/Dsred^-^ cells. Luminal CD44^lo^ or CD44^hi^ cancer cells obtained from MCF7 cells were co-cultured in vitro. The collective expansion was observed by time-lapse microscopy. Scale bars, 50μm. (E)Heat maps of unsupervised hierarchical clustering data are shown depicting differentially expressed genes in EpCAM^+^/CD45^-^/CD44^hi^ and EpCAM^+^/CD45^-^/CD44^lo^ subsets (blue and yellow indicate higher and lower expression, respectively) with fold changes ≥2 and FDR≤0.05. (F) Differential gene expression verified by quantitative RT-PCR in FACS-purified CD44^hi^ and CD44^lo^ subpopulations from primary breast cancer cells. GAPDH was used as a loading control. (G) Confocal images of localization of E-cadherin, β-catenin, ZO-1, and F-actin at the leading edge and centre of the migrating clusters. Data are representative of at least three experiments. Scale bar, 25 μm.

To further determine how CD44^lo^ and CD44^hi^ cells act in vivo, CD44^hi^ and CD44^lo^ cells sorted from MMTV tumors were immediately injected orthotopically into NOD/SCID mice at serial dilutions from 200 cells to 5×10^5^ cell. In most cases, CD44^hi^ cells generated 3- to 23-fold larger tumors compared to their CD44^lo^ counterparts and presented greater tumor-initiating ability (Fig. 2E). However, we observed that the CD44^lo^ cells formed tumors even in relatively low numbers of cells, although with a smaller size and longer duration at causing tumors (Fig. 2E). These results illustrated that CD44^hi^ cells arising in vivo behave much like the CSCs in that they exhibit higher tumor-initiating and tumor growth potential than their CD44^lo^ counterparts. As CD44 is widely accepted as a CSC marker, the results raised the question of how CD44^lo^ populations that apparently lacked of CSC features were able to generate tumors when transplanted into mice.

### 3. CD44^hi^ cells led collective invasion of luminal breast cancer cells

To observe the migration of CD44^hi^ cells and CD44^lo^ cells, we cultured two confluent epithelial monolayers within two confined wells separately and then remove the confinement, triggering cluster migration in the forward facing direction(Fig.3A) as modified from previous report(Das et al, 2015). The CD44^hi/DsRed+^ and CD44^lo/DsRed-^ cells were isolated from MCF7/naïve cells and MCF7/DsRed cells by FACS. The migration of CD44^hi^ cells and CD44^lo^ cells was monitored during collective migration using time-lapse microscopy. Result showed that the migration rate of CD44^hi^ cells is significantly higher than that of CD44^lo^ cells(Fig.3B; Supplemental Movie 1-2). Knockdown and overexpression experiments showed that CD44 promoted collective migration, with CD44 depletion expectedly inhibited the migration of CD44^hi^ subpopulation and CD44 overexpression encouraged the motility of CD44^lo^ subpopulation (Fig.S2).

Moreover, to investigate whether CD44^hi^ subset leads the migration of CD44^lo^ subset and thereby promotes collective invasion, confocal time-lapse imaging of the cocultures of CD44^hi/DsRed+^ and CD44^lo/DsRed-^ subsets was performed. Imaging result showed that the leading cell is always a CD44^hi^ subset and that the CD44^lo^ cells move behind (Fig.3, C and D; Supplemental Movie 3-5).

### 4. The molecular phenotype was distinct between the CD44^hi^ and CD44^lo^ cancer cells generated during the collective invasion

In order to gain further insights into the molecular mechanisms enabling the observed differences between the two cell subsets in breast cancer, gene-expression signatures of FACS-sorted epithelial cells were analyzed. CD44^hi^ cells that had been grown as leader cells or CD44^lo^ cell as follower cells from the primary breast tumors were collected. Then, RNA sequencing was conducted for detecting the acquired gene expression at the CD44^hi^ leader cells and compared with other CD44^lo^ follower cells. As a result, 3,826 genes were differentially expressed between the two subsets (R2-fold; p < 0.05) (Fig.S3A, B). Unsupervised hierarchical clustering of differentially expressed genes was involved in three major pathways: focal adhesion, EMT and RAS signaling pathway.

To address the factors underlying the different phenotypes, we analyzed the most significantly changed genes bioinformatically. Gene Ontology (GO) and KEGG functional enrichment analysis revealed 10 distinct terms enriched in the CD44^hi^ cells (Fig. S3C). According to the high invasiveness in CD44^hi^ cells, several genes related to cell motility and modifiers of the extracellular microenvironment were among the most differentially expressed genes, with lower levels in the CD44^lo^ cells, including MMP2, MMP9, Plau, CLDN-1, −3, −11, and −7. Moreover, the CD44^hi^ cells expressed a number of mesenchymal markers not present in CD44^lo^ cells (Fig. 3E), including reduction of epithelial markers including CDH1 (E-cadherin), JUP (γ-catenin), krt8, and krt18 and increase of mesenchymal markers including FN1 (fibronectin), Vim (vimentin), COX-2(Ptgs2), and ZEB2. This was confirmed by quantitative real-time PCR (qRT-PCR) (Fig. 3F). However, how the CD44-associated invasive gene set revealed in CD44^hi^ subset coordinates with each other and contributes to collective invasion are unknown.

Next, we investigated the localization of some additional mesenchymal markers that might explain the sustained invasion of carcinoma cells (Fig.3G). Surprisingly, besides the difference of E-cadherin RNA expression between CD44^hi^ and CD44^lo^ cells, epithelial markers E-cadherin and ZO-1 are missing at the lamellipodia in leading edge cells but remaining_membrane-localized in the rear end of leading cells at the contacting area with the follower cells, with a similar distribution of β-catenin and a reorganization of F-actin(Fig.3G, Fig. S4). The migrating leader cells displayed branched F-actin filaments at the leading edge cells with a concomitant forward-moving lamellipodia. Together, our results suggest that a complete EMT program is not achieved in the invading leader cells, despite vigorous invasiveness of cells into the ECM.

Although tumor organoids culture-based studies have described a critical role for K14+ and p63+ cells in leading tumor cells during collective invasion(Cheung et al, 2013), we did not observe any differences in the expression of K14 or p63 gene between CD44^hi^ MMTV-PyMT and CD44^lo^ MMTV-PyMT primary breast tumor cells. Besides tumor cells, tumor organoids in these studies contain tumor cells and stromal cells too, which were located at the boundary of tumor-stromal interface(Corsa et al, 2016) and influenced cell invasiveness. What’s different here is that we obtained CD44^hi^ and CD44^lo^ cell by FACS depending on EpCAM^+^/CD45^-^, in which stromal cells were excluded. Moreover, observations of luminal cell lines also revealed that CD44 was located at the migration edge and tumor-stromal boundary of xenografts (Fig.1C).

### 5. The CD44^lo^ and CD44^hi^ states can spontaneously switch to each other

To understand why the CD44^lo^ subsets at lower cell numbers including 2000cells/site were able to generate tumors when xenografted into mice, we investigated the dynamic changes of CD44 in the sorted CD44-different subpopulations. In the purified CD44^lo^ group, the proportion of CD44^hi^ cells was found to increase with cell expansion (Fig.4A). Alternatively, in the purified CD44^hi^ group, the proportion of CD44^lo^ cells was also expectedly found to increase with collective cell growth. Significantly, over a period of time, the level of CD44 in purified CD44^hi^ and CD44^lo^ group was stabilized to the level typical of the unsorted parent cells, indicating that the two cell subsets can shift from CD44^hi^ phenotype to the CD44^lo^ state in a dynamic equilibrium that maintains the proportion of CD44^hi^ cells located at the migrating tips (Fig.4A). The failure of the sorted CD44-different subsets to be stably propagated indicated that mesenchymal-like CD44^hi^ leading cells do not represent a stable state within an individual tumor.

**Fig. 4.**
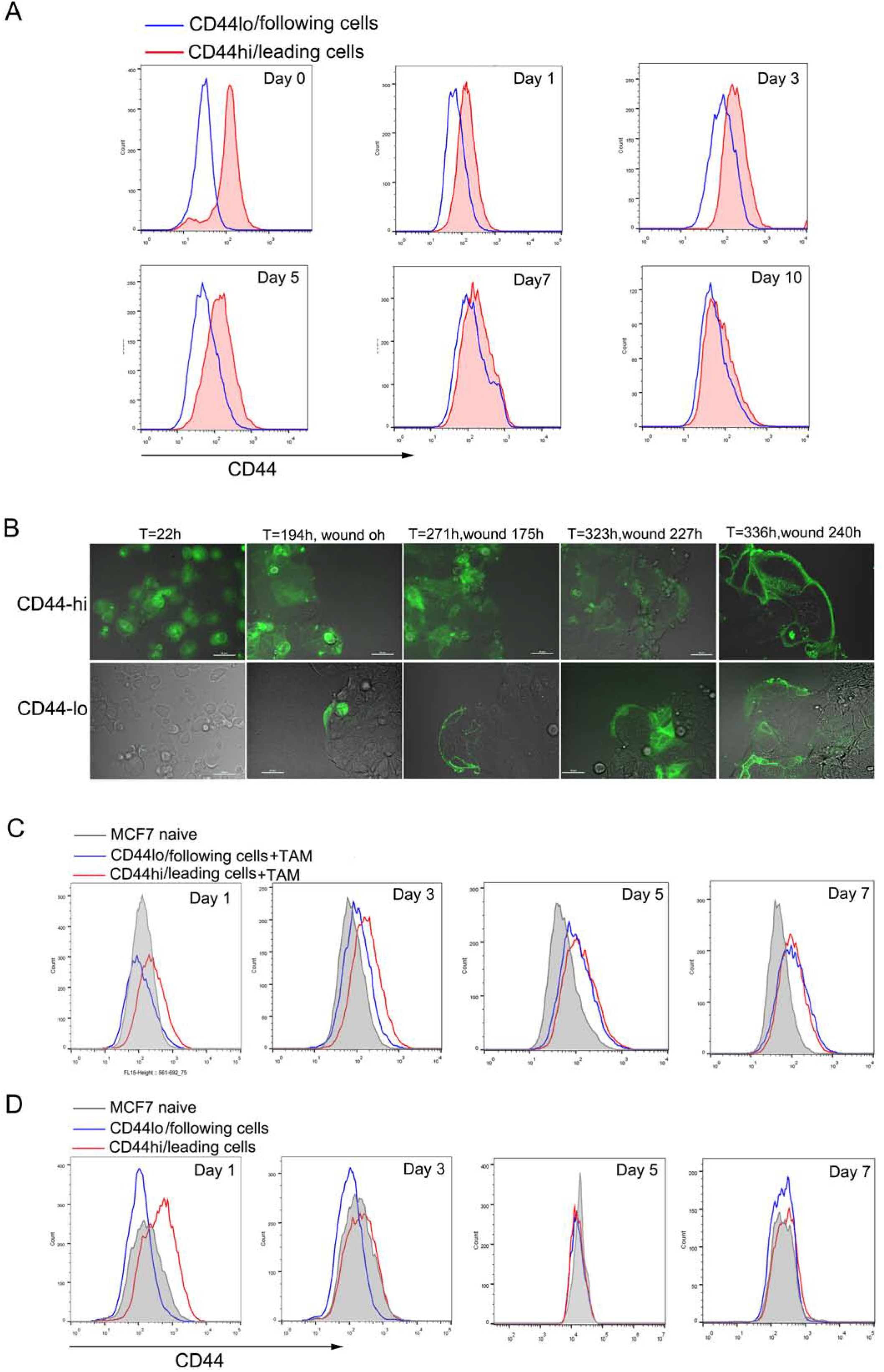
Inducible shift of CD44 expression occurs during collective invasion. (A) Flow cytometric analysis of the changes of CD44 expression in CD44^hi^ and CD44^lo^ subpopulations during cell expansion in vitro. Purified luminal CD44^lo^ or CD44^hi^ MCF7 cells were cultured in vitro and analyzed by FACS for proportion of CD44^lo^ and CD44^hi^ components. Luminal BrCa cells may convert directly from CD44^hi^ to CD44^lo^ states and vice versa. (B) Time-lapse microscopy of CD44 switching in CD44hi and CD44lo fractions sorted from MCF7 cells, expressing YFP under the control of the CD44 promoter (CD44 promoter-YFP). (C) Flow cytometric analysis of changes of CD44 levels after stimulated with TAM. TAM cells were obtained after THP-1 cells were stimulated to have M2-phenotypes. Purified CD44^hi^ or CD44^lo^ cells obtained from MCF7 cells were incubated in the absence or presence of TAM cells for 0-7days using a Transwell coculture system, then examined for CD44 levels by FACS. (D) Flow cytometric analysis of changes of CD44 levels after treatment with hypoxia. Purified CD44^hi^ or CD44^lo^ MCF7 cells were incubated in hypoxia and subjected to flow cytometric analysis. Scale bars, 50μm.

Alternatively, to observe the CD44-dependent switching directly, we generated MCF7 cells carrying a CD44 promoter biosensor in which YFP reports on CD44 gene expression. In these cancer cells, CD44^lo/YFP+^ and CD44^hi/YFP-^ subpopulations were sorted and cultured in vitro, separately. We observed the transition from CD44^lo^ to CD44^hi^ states of individual cells at the leading edge and the switch to CD44^lo/YFP+^ states in the CD44^hi/YFP-^ cells derived follower cells (Fig.4B, Supplemental Movie 6-7). Following phenotypic conversion, the newly generated CD44^hi/YFP-^ cells dynamically extended and retracted membrane protrusions and led migration of trailing CD44^lo/YFP+^ cells (Fig.4B, Supplemental Movie 6-7). Taken together, these data informed that CD44^lo^ cells were nonetheless able to generate new CD44^hi^ cells, albeit at a low frequency. The invasive CD44^hi^ leader phenotype may convert directly from CD44^hi^ to CD44^lo^ states along with the collective cell migrating, not a stable lineage.

### 6. Microenvironmental stimuli enhanced the rate of interconversions between CD44^lo^ and CD44^hi^ states

Extensive evidence indicated that the stromal microenvironmental signals could influence cancer cell plasticity and tumorigenicity(Friedl & Alexander, 2011). Among these signals, hypoxia and tumor associated microphage (TAM) stimulation have been shown to potently activate EMT program and cancer stemness(Chaffer et al, 2013; Gregory et al, 2011). Accordingly, we examined whether hypoxia or TAM stimulation could affect the spontaneous transitions between CD44lo and CD44^hi^ states of luminal BrCa cells. Purified CD44^lo^ and CD44^hi^ epithelial cells were cultured under hypoxic condition or cocultured with TAMs, respectively. TAMs cells were obtained after THP-1 polarized to M2-phenotype macrophages, according to our previous report(Zhang et al, 2016). Results showed that the treatment of hypoxia to CD44^lo^ cells results in a rapid generation of a CD44^hi^ subpopulation, and similar results occur when isolated CD44^lo^ cells are cocultured with TAMs(Fig. 4C). Conversely, CD44 expression was reduced in CD44^hi^ cells treated with hypoxia (Fig. 4D). Additionally, coculture with TAMs results in the rapid increase of CD44 levels in both of isolated CD44^lo^ cells and CD44^hi^ cells (Fig. 4C). The results suggested that tumor microenvironment contributes to the formation of CD44^hi^-leader cells in the leading front. Moreover, breast cancer microenvironment has an enhanced inflammatory network, in which IL-6 is reported to be an important regulator(Yadav et al, 2011). However, when rhIL6 is added to the medium, the conversion between CD44^lo^ and CD44^hi^ was not influenced, suggesting that secreted IL6 is not indispensable for generating CD44^hi^ leader cells from epithelial tumor (Fig. S5). Together, these data indicated that microenvironmental signals in individual tumor enhance the rate of interconversions between CD44^lo^ and CD44^hi^ subsets in luminal breast cancer and contribute to the generation of CD44^hi^-leader cells.

### 7. The expression patterns of CD44 in the invasive CD44^hi^ leading-cells and the CD44^lo^ following-cells

To further identify the molecular mechanisms responsible for the switch of CD44, we investigated the expression pattern of CD44 variants. This pattern was detected in a reverse transcription polymerase chain reaction (RT-PCR) assay by a primer pair spanning the entire variant region(Banky et al, 2012; Yang et al, 2015). Our results showed that there is no difference of CD44s expression between CD44^hi^ and CD44^lo^ cells, whereas, the mRNA levels of CD44 variants were higher in the CD44^hi^ cells compared with those CD44^lo^ cells (Fig.5, A and C). When co-cultured with TAMs, expression of the CD44v was strongly up-regulated in both of the CD44^hi^ and CD44^lo^ subsets (Fig.5, B and D). Our results revealed that CD44 alternative splicing is differentially regulated during collective invasion, resulting in a switch in expression from the predominant isoform, CD44s, to the enriched variable exon-containing CD44v isoforms.

**Fig. 5.**
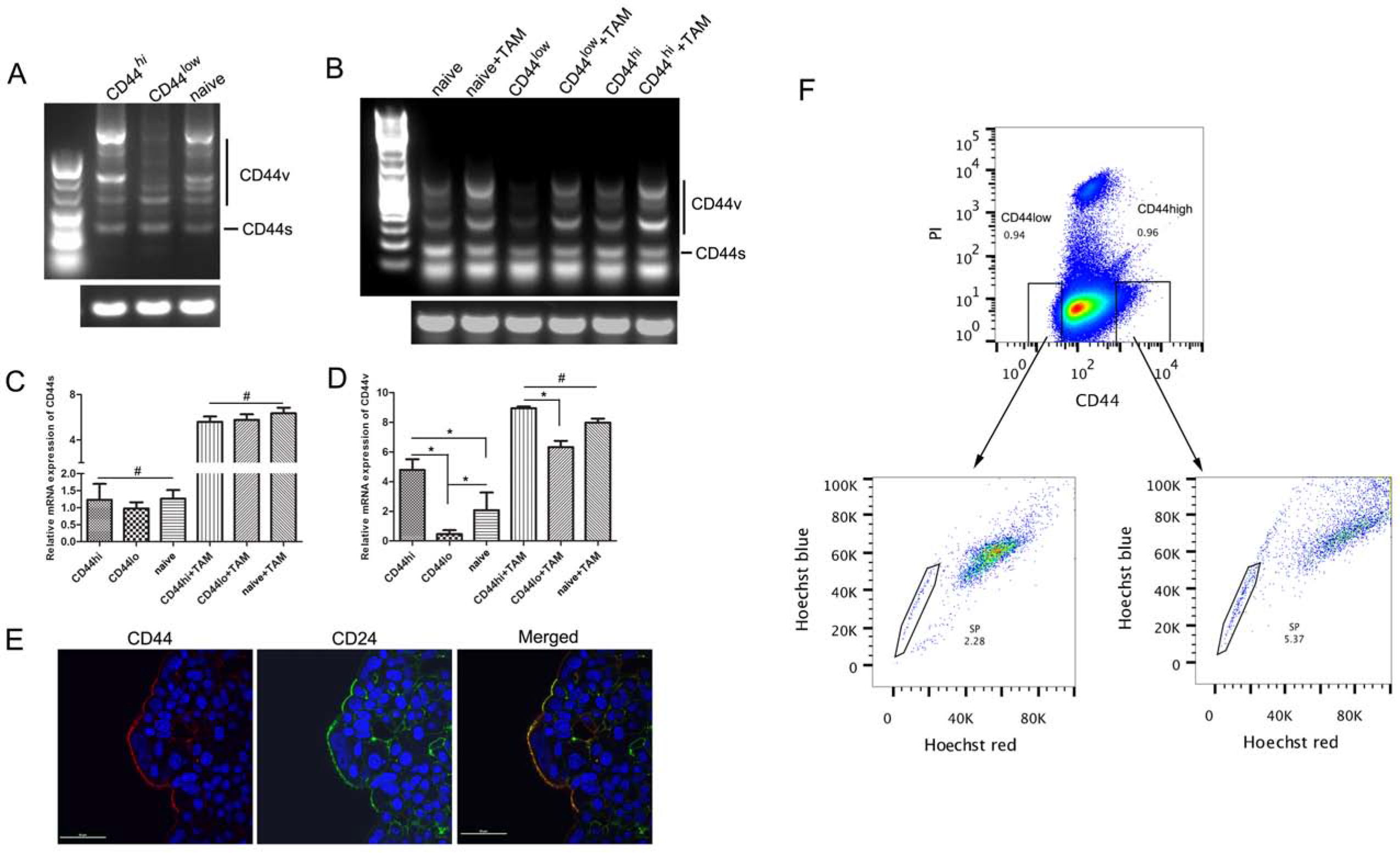
CD44 expression pattern with CD44-dependent phenotype and characterization of CSC-like cells during collective migration. (A) Reverse transcription PCR (RT-PCR) analysis of CD44 expression pattern in the CD44^lo^ or CD44^hi^ subpopulation generated from collective migration. Using a primer pair spanning the entire variant region, the CD44 variants expression was detected. (B) CD44v expression was boosted by TAM stimulation. Co-culture system of TAM cells and FACS-purified CD44^lo^ or CD44^hi^ MCF7 cells were used as described before. After co-incubation for 3 days, cells were harvested and analyzed by RT-PCR for the expression pattern of CD44. (C) Quantitative analysis of CD44s in the CD44^lo^ or CD44^hi^ subpopulation determined in (A) and its changes induced by TAM treatment in (B).(D) Quantitative analysis of CD44v in cell subpopulations before and after TAM stimulation in (B). Relative expression levels of CD44s or CD44v were normalized to GAPDH. Error bars indicate SD; n = 4. (E) Immunocytochemistry analysis of the expression of cell surface CD44 and CD24 in collective migrating cell clusters. MCF7 cells were cultured in the wound chamber (ibid). After a confluent monolayer was formed, the culture-insert was removed to trigger a sheet migration of MCF7 cells. Then, cells were fixed for immunofluorescence assay at 72 h, after the wound was made. Representative images were taken under a confocal microscope. Scale bars, 100 μm. (F) Hoechst 33342 efflux assay of SP and non-SP cells in CD44^hi^ and CD44^lo^ subpopulations. CD44^hi^ or CD44^lo^ cancer cells (upper) were separately incubated with Hoechst 33342 dye and then analyzed by flow cytometry.

### 8. Changes of CSC population during the CD44 switching

It is widely accepted that CSCs are contained exclusively in the CD44^hi^ cancer cells(Al-Hajj et al, 2003; Visvader & Lindeman, 2012). Therefore, we observed the localizaton of CSC cells in migrating cohesive clusters of luminal cancer cells using the expression of CD44 and CD24. Results showed that only a small CD44^+^/CD24^-^ subclone was detected in migrating front cells, but not all of leading cells were in CD44^+^/CD24^-^ state (Fig.5E). Whereas, a subpopulation of cancer cells with CD44^+^/CD24^+^ positive accounted for the dominant population of the leading cells. The result illustrated that the nature of CD44^hi^/leading cells, residing in the migrating tips that produce higher tumor-initiating and tumor motility potential than their CD44^lo^ follower counterparts, is not depending on the CSC-like characteristics.

Furthermore, to test whether the changes of CD44 levels on its own would allow enrichment of CSCs from BrCa cells, we analyzed the changes of CSCs proportion during CD44 switching in collective expansion of luminal BrCa using the Hoechst 33342 dye exclusion method (side population: SP). Accordingly, the proportion of CSC in luminal CD44^hi^ or CD44^lo^ cancer cells was measured by flow cytometry (FCM). We found that luminal CD44^hi^ subpopulations contained CSC cells with a proportion of CSC cells (less than 5.5%). Meanwhile, luminal CD44^lo^ subsets also comprised a small proportion of CSC cells (less than 2.2%) (Fig.5F). This result implied that the switching of CD44 might not be corresponding to conversions of the stemness properties of non-CSCs-to-CSCs.

### 9. Dynamic colocalization of merlin/ezrin and CD44 enabled collective migration

Of known intracellular CD44 binding partners, ezrin and merlin were reported important in contractile transduction and assembly of cortical actin during cell migration(Fehon et al, 2010). The distribution change of ezrin or merlin in response to various signals could control the lateral cell mobility(Chiasson-MacKenzie et al, 2015). We then investigate how the binding partners are involved in enabling the observed collective cell movements. A confluent epithelial BrCa monolayer was cultured to generate sheet migration by scratch-wounding. In the early time points of the sheet migration, the expression of ezrin was very low at the wound edge, which was coinciding with that of CD44 (Fig.6 A, left, Fig.S5). In contrast, after 72h of migration, eztin was enriched in the leading edge cells, displaying co-localization with increased CD44 and suggesting complex formation between ezrin and CD44(Fig.6A, right, Fig.S6). Therefore, the increased CD44-Ezrin complexes at the leader cells presumably stabilize directional cell motility. Simultaneously, within the first few minutes of wound, merlin mostly presented at cohesive cell-cell boundaries (Fig.6B, left, Fig.S6). Interestingly, after collectively migrating for 72h, in the leading cells, merlin molecules transferred to the cytoplasm, whereas remains stable in stationary follower cells (Fig. 6B, right, Fig.S6). The spatial relocalization of merlin in the cell-sheets suggested that the migration-induced cellular pulling-force triggered by leader cells resulted in a release of cortical merlin. Merlin is known to control lamellipodium formation(Das et al, 2015). The spatial location of merlin and formation of CD44-Ezrin complexes, accompanied with CD44 switch in the moving cell-sheets, presumably stabilizes the formation of invasive lamellipodium. Merlin/ezrin and CD44 might form a molecular switch that supports collective cell migration.

**Fig. 6.**
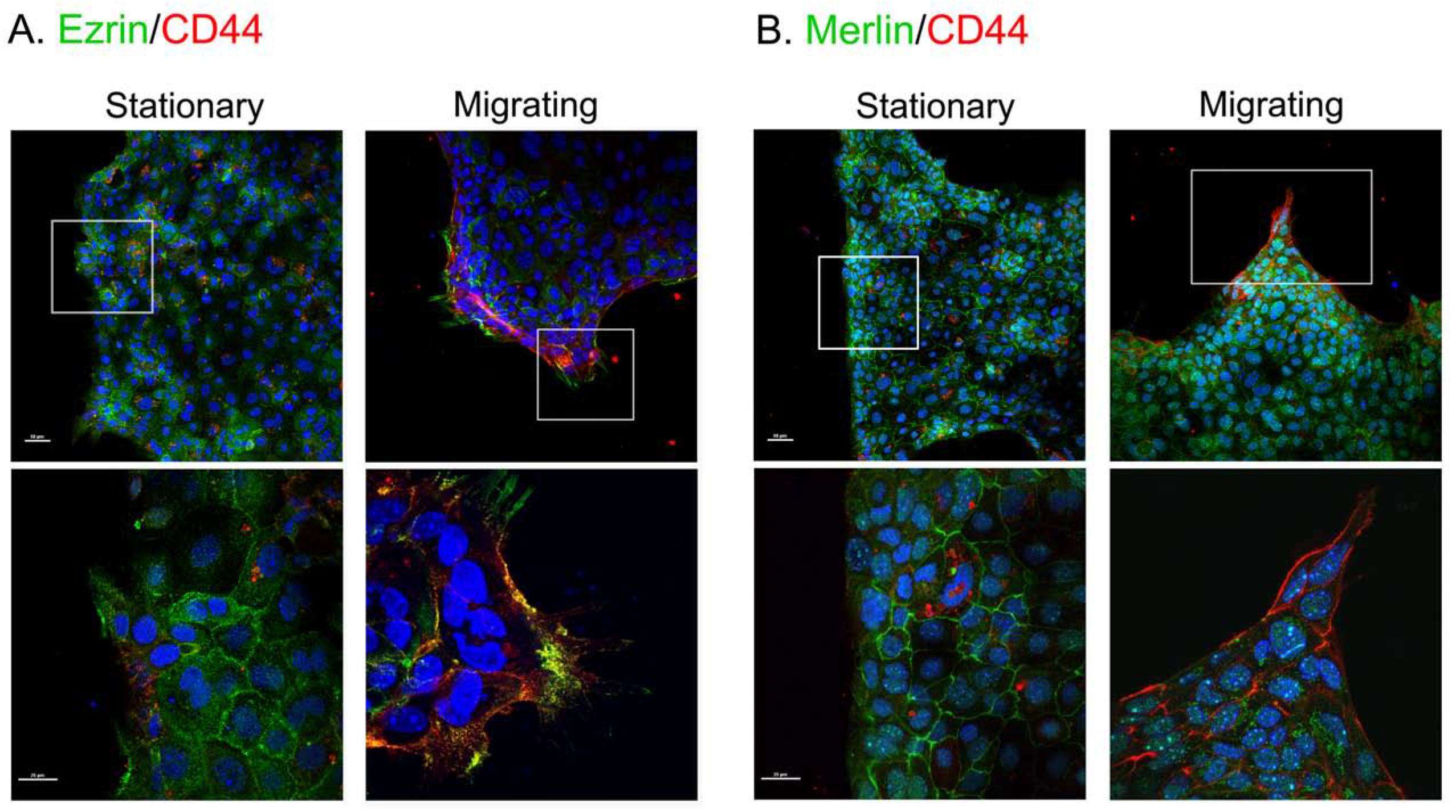
Co-localization of CD44 and Merlin or Ezrin in stationary and migrating cells during collective migration. (A) Co-localization of CD44 and Ezrin in stationary (left) and moving (right) monolayer. EpCAM^+^/CD45^-^/CD44^hi^ or EpCAM^+^/CD45^-^CD44^lo^ cancer cells were purified from MMTV/PyMT mice, and then cultured in the wound chamber (ibid). A sheet migration of epithelial cells was generated as described before. Then, cells were fixed for Ezrin and CD44 expression at 0h and 24h post wounding by immunofluorescence assay. Representative images were presented. Scale bars, 50 μm. (B) Localization of CD44 and merlin in stationary (left) and migrating (right) monolayer. A sheet migration of epithelial cells was made as previously described. Merlin and CD44 expression at 0h and 24h post wounding were determined by immunofluorescence assay. Scale bars, 50μm.

### 10. The shift of CD44^hi^ and CD44^lo^ states within tumors in mouse xenografts

To further address the reciprocal conversion between sorted CD44^hi^ and CD44^lo^ subsets, we performed a mixed xenograft experiment (Fig.7A) in SCID mice involving co-injection of purified CD44^hi^ and CD44^lo^ cells in a mixed population from MMTV-PyMT mice. Initial cells from purified CD44^hi^ cancer cells, CD44^lo^ cancer cells, or the mixture group were xenografted orthotropically, separately. Accordingly, tumor size was measured with calipers. The tumors that arose from different subpopulations as described above were digested and analyzed for CD44 expression by flow cytometry (FCM). Results showed that CD44^lo^ BrCa cells displayed relative weaker tumor-initiating ability, with the smallest tumors among the three groups (Fig.7B). However, tumors derived from each subpopulation comprised almost the same proportion of EpCAM^+^/CD45^-^/CD44^lo^ (p=0.4068) and EpCAM^+^/CD45^-^/CD44^hi^ cells (p=0.4891) (Fig.7, C and D). This suggested that luminal CD44^lo^ BrCa cells generate new CD44^hi^ cells in vivo, therefore initiating tumor formation, although at a low speed. In comparison with tumors generated by CD44^lo^ subset alone, the tumors derived from coinjection group were obviously larger. Interestingly, although only half the amount of CD44^hi^ cells is used in the mixed group (Fig.7E), in comparison with tumors generated by CD44^hi^ subset alone, there is no obvious difference in tumor size between the CD44^hi^-derived tumors and the mixed group-derived tumors. These findings indicate that luminal CD44^hi^ to CD44^lo^ conversion happened in vivo so that they achieve final equilibrium state of CD44 levels.

**Fig. 7.**
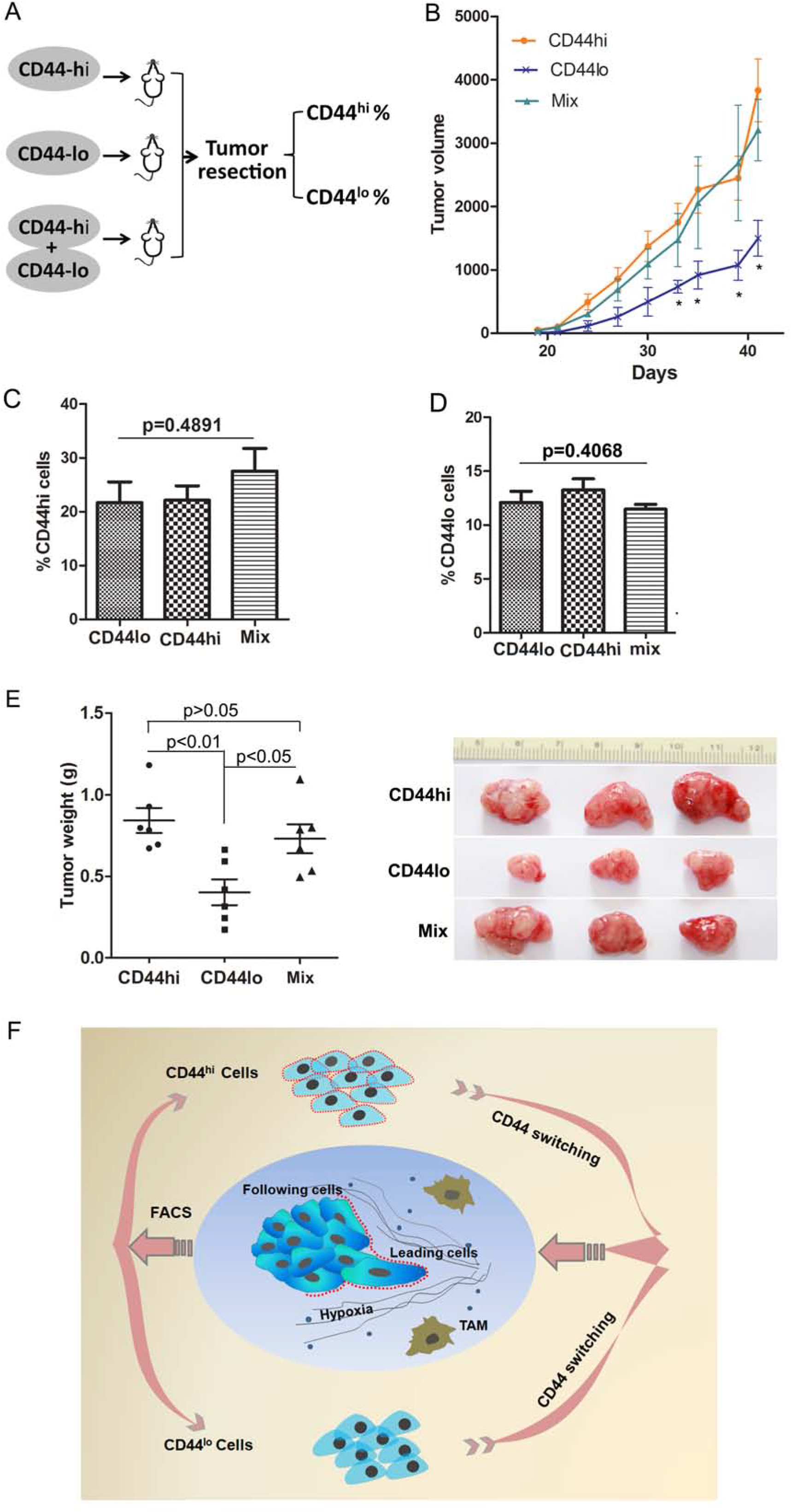
Conversion of CD44^hi^ and CD44μ cells within tumors in mouse xenografts. (A) Scheme of mixed xenograft experiment. The CD44^hi^ cells were combined (or not) with CD44^lo^ cells from MCF7, injected into NSG-SCID mice.(B) Tumorigenicity of FACS-sorted luminal-like CD44^lo^ or CD44^hi^ BrCa cells following orthotopically injected into SCID mice (n≥6/group).(C, D) Quantification of CD44^hi^ or CD44^lo^ cells generated from purified-CD44lo-, CD44hi-, or mixed group-derived tumors in (A). Cells were obtained from excised tumors 21 d after injection and analyzed by FCM.(E) Weights of tumors generated from luminal CD44^lo^, CD44^hi^, or mixed cell populations and representative tumor image.(F) Scheme of CD44 switching in leading collective cancer cell invasion. A collective invasion model shows phenotypic interconversion between two subpopulations, a highly invasive CD44^hi^/leading cell (red membrane) at the front of the invading cell clusters and CD44^lo^/following cells frequently occurred during collective invasion, which is spontaneous and sensitive to tumor microenvironment.

## Discussion

Clinically, luminal breast cancers can collectively invade as cohesive cell clusters, with about 20% of cases that eventually undergo metastasis(Kennecke et al, 2010). The molecular basis of the transition from in situ to invasive state in breast cancer remained elusive till now. Recently, the plasticity and continuous evolving of cell populations within the same tumor bring more challenges for explanation of the mechanisms involved(Friedl & Alexander, 2011).

### CD44^lo^-to-CD44^hi^ Conversion Generates Invasive Leader Cells

The present work reveals that CD44^hi^ cancer cells are the leading subpopulation in collective invading luminal cancer, which could efficiently lead the collective invasion of CD44^lo^/follower cells. Indeed, analysis of the gene expression signature of CD44^hi^/leading subpopulation identified a cohort of hybrid epithelial/mesenchymal genes and key functional co-regulators of collective invasion, which was distinct from CD44^lo^/following cells. Furthermore, luminal CD44^hi^ subpopulation was readily switching to CD44^lo^ state along with collective cell movements, which is spontaneous and also sensitive to tumor microenvironment. Meanwhile, converting to CD44^hi^ state was presented in luminal CD44^lo^ subsets. Thus, the CD44^hi^ leader cells featuring an intermediate EMT are not a stable subpopulation in breast tumor (Fig.7F). The ability of epithelial cancer cells to acquire an invasive mesenchymal-like phenotype associated with the CD44 switching from a CD44^lo^ to a CD44^hi^ state, accompanied with a shift of CD44s-to-CD44v, suggests that invasion, metastasis, and patient outcome may be influenced by the inducible interconversions between CD44^lo^ and CD44^hi^ state. Collectively, the transition from in situ to invasive behavior of luminal-type BrCas could be considered to be due to this plasticity and ability to generate CD44^hi^ carcinoma cells with enhanced invasion-initiating powers.

### CD44 Is Spatially Distributed In Invasive Cell Clusters

CD44 is revealed to contribute to promote tumor progress by cooperating with coreceptors(Godar et al, 2008; Ponta et al, 2003). In our study, we found that the differences of CD44 levels appear to be closely related to the differences in cell molecular subtypes, with generally lower level in luminal BrCa than in basal BrCa, corresponding to its less aggressive and better prognosis than basal BrCa in clinical cases(Roswall et al, 2018). Additionally, in luminal BrCas, CD44 was enriched at the leading front of the invading cell strands, whereas in the inner region of the collective clusters it was maintained at a lower level. The CD44^hi^ cells residing have a strong impact on the overall invasive behavior of these cohesive tumor cells. Accordingly, as a tumor-promoting protein(Ahmed et al, 2016) and a useful CSC marker(Al-Hajj et al, 2003; Prince et al, 2007), the spatial distribution of CD44 in invasive cell clusters, appears to be responsible for the profound differences in cell invasive ability among heterogeneous subpopulations.

### CD44^lo^-to-CD44^hi^ Conversion Is Accompanied by Switch of Molecular Phenotypes

Our further study showed that the invasive phenotype of CD44^hi^/leading cells is obtainable and dependent upon the CD44^lo^-to-CD44^hi^ conversion, which is in line with the notion that cancer cells may undergo reversible switching of phenotype under deliberate cell regulating program. As suggested before, switch of phenotypes between subpopulations modulates the behavior of cancer cells in solid tumors(Roswall et al, 2018). In particular, there have been multiple observations about phenotype switching between subpopulations of breast cancer cells, that caused chemotherapy failure and tumor metastatic dissemination(Hanahan & Coussens, 2012; Yu et al, 2013). Evidences revealed that CSC-like cells may arise from non-CSCs and vice versa, and the molecular subtype of breast cancer can convert from basal-like phenotype to luminal-like state, which may influence the sensitivity to chemotherapy (Iliopoulos et al, 2011; Yang et al, 2012). In this study, results indicate that the partial-mesenchymal CD44^hi^ subpopulations contain a small subclone of CSC cells; however, the following cells also have a small subset of CSC-like cells. Therefore, the CD44 switching is not equivalent to the shift of non-CSC-to-CSC state. Whether traits attributed to CSC cells in previous studies, such as resistance to chemotherapy, are conferred to CD44^hi^/leading cells or CD44^lo^/following cells remain to be determined. However, although we cannot rule out the possibility that CSCs may play a critical role in attaining invasive phenotype of leader cells, we demonstrated that CD44 switching plays a critical role in driving formation of leading cells.

Furthermore, our results showed that the isoform switch from CD44s to CD44v occurred during the collective invasion. Knowledge about the precise function of CD44s and CD44v is as yet incomplete, rather controversial. It is reported that CD44v is related to metastatic properties whereas CD44s confers nonmetastatic properties(Morath et al, 2018; Yae et al, 2012). CD44v isoforms are also associated with stem-like properties in many different cancers(Banky et al, 2012; Guo & Frenette, 2014; Ishimoto et al, 2011). However, cancer cells with CD44s were showed to be more invasive and stronger drug resistance in pancreatic cancers(Zhao et al, 2016). The isoform switching from CD44v to CD44s was reported to contribute to EMT and breast cancer progression in immortalized normal cells where CD44s, not CD44v, has been manifested to be crucial for successful complete EMT(Brown et al, 2011). Taken together, although being disputative, the isoform switching of CD44 mediated by alternative splicing has important impact on malignant behaviors of cancer cells. Our results thus revealed that CD44 alternative splicing happens in response to the stimuli of TAMs, resulting in a shift in expression from CD44s to CD44v isoforms. Through isoform switching, the obtained high levels of CD44 isoforms might be responsible for the distinct cellular functions of cell subpopulations, including invasion and tumor-initiating ability.

### Leading Cells Have CD44-Dependent Invasion Program

We observed large, CD44-dependent differences between CD44^hi^ and CD44^lo^ subpopulations in the same luminal breast tumor. In accordance with the ability of CD44^hi^ cancer cells in driving collective invasion, we detected a specific molecular program in CD44^hi^ subpopulation. Suppression of epithelial markers (CDH1 and JUP) and upregulation of mesenchymal markers (Vim, FN1, Ptgs2, and ZEB2) occurred in the CD44^hi^/leading cells. These changes are reported to be sufficient to activate complete EMT(Drees et al, 2005). However, at the same time, upregulation of tight junction proteins claudin 1, 3, 4 and 7 was also observed in the leader cells, which is opposite to the classic EMT program(Ribeiro & Paredes, 2014). Moreover, krt8 and krt18, typically expressed in epithelial cells, were reduced in CD44^hi^/leader cells compared with that in CD44^lo^/follower cells. Thus, the CD44^hi^/leading cells display a hybrid phenotype with both epithelial and mesenchymal features, implying that it did not undergo a complete EMT; instead, undergo partial EMT, an intermediate state. In agreement with this result, loss of krt8/krt18 in epithelial cancer cells was shown to increase cancer invasiveness without inducing EMT(Fortier et al, 2013). The hybrid E/M state has been described as a metastable phenotype(Lee et al, 2006), generating more plastic and metastable cancer cells(Klymkowsky & Savagner, 2009). Therefore, all these alterations might improve the formation of invasive tips, leading to an increase in cell motility and invasion capacity, and orchestrate the collective movement. Although expression of CD44 is reported to be regulated by p53, p63(Godar et al, 2008), and ESPR1/2(Brown et al, 2011), we did not observe any difference of those genes expression during the switch of CD44 in luminal BrCa. The precise mechanisms related need further exploration.

In addition, MMP2, MMP9 and Plau expression were increased in CD44^hi^/leader cells, in comparison with CD44^lo^/follower cells, which were demonstrated to accelerate invasion through remodeling of the ECM. Furthermore, analysis on the distribution features showed that CD44^hi^/leading cells presenting specific characteristics with some epithelial markers (E-cadherin, β-catenin, and ZO-1) were missed at the leading edge of lamellipodia, but remained at the contacting area with the following cells. Taken together, our data suggest that leader cells could possess all the gene expression required for enhanced cell invasion capacity but remain cell-cell junction with the follower cells. All these hybrid traits exhibited a transient EMT state, with an increased plasticity and invasiveness, which might be an additional explaination why EMT is dispensible for metastsis(Zheng et al, 2015).

### Merlin/Ezrin and CD44 Coordinately Regulate Collective Invasion

Observations have suggested that, through forming coreceptor complexes with various receptors and signaling kinases, CD44 drives numerous signaling pathways in tumor-promoting effects (Godar et al, 2008) (Ponta et al, 2003). Also, CD44 serves as a picket attaching the actin cytoskeleton to the plasma membrane, connecting cyto- and pericellular signaling to modulate receptors mobility(Freeman et al, 2018). In this study, confocal microscopy studies confirmed the co-localization of enriched CD44 and ezrin at the front ends of a migrating leader cell, while merlin relocalizing to the cytoplasm. Meanwhile, in the follower cells, merlin mostly concentrated at cell-cell contact sites together with E-cadherin and β-catenin. Ezrin was revealed to interact with both of CD44 and F-actin, forming a bridge between CD44 and the actin cytoskeleton, mediating changes in cell morphology and migration(Donatello et al, 2012). Merlin was reported to act as a mechanochemical transducer, coordinating lamellipodium formation and collective movement of cell clusters during wound healing(Das et al, 2015). Therefore, the reciprocity of Ezrin-merlin-CD44 association in migrating cell clusters may help maintain collective cell movement during breast cancer cell invasion. However, further studies are needed to clarify the mechanism.

## Conclusion

In summary, the study indicated the potential values of CD44 switching and shifts of CD44s/v in explaining the mechanism of metastasis and predicting the progression from luminal breast cancers in situ to invasive carcinoma. This inducible acquirement of CD44 cast some challenges on the molecular targeting therapy, as has been demonstrated in non-CSC and CSC conversions. In addition, spatial localization of distinct phenotypic cells in solid tumors might lead to the generation of distinct niches around the invading cell clusters that favor the cohesive cell invasion as a whole. Therefore, in the future, a deeper understanding of the intratumor plasticity and their consequences might provide new insights for the mechanisms underlying collective invasion and metastasis.

## Experimental Procedures

### 1. Antibodies and Immunofluorescence

For immunofluorescence, cells were fixed in 4% paraformaldehyde, permeabilized with Triton-X-100/PBS, and blocked in 1×PBS + 1% BSA. Primary antibodies were incubated overnight at 4°C, 1× PBS + 1% BSA. Secondary antibodies with Alexa Fluor 488 or 647 (Abcam) were incubated 1 h at room temperature. Images were analyzed under confocal microscopy (Nikon A1, Tokyo, Japan).Primary antibodies were CD44 (Abcam, ab119348), E-cadherin (Abcam, ab40772), β-catenin (Abcam, ab32572), Merlin (Bioworld Technology, BS3663), Ezrin (Cell Signal, 3145s), and phalloidin (Abcam, ab143533).

### 2. Cell culture and Mice

Human breast cancer cell lines MCF-7, T47D, and ZR-75-30 were bought from American Type Culture Collection (ATCC) and cultured in MEM, RPMI-1640 medium, or Dulbecco’s Modified Eagle Medium (DMEM) supplemented with 10% fetal calf serum, 100 U/ml penicillin, and 100 mg/ml streptomycin. All cells were grown to 85% confluency for experiments.

In this report, we explore the role of CD44 in tumor progression using the mouse mammary tumor virus LTR-driven polyoma middle T antigen (MMTV-PyMT) transgenic mouse, which has been proved to be a reliable model of human luminal breast cancer (Guy et al., 1992; Herschkowitz et al., 2007; Lin et al., 2003). MMTV-PyMT (FVB/n) mice were from Jackson Labs. All protocols involving mice were evaluated and approved by our Institutional Animal Care and Use Committee and performed under veterinary supervision.

### 3. Immunohistochemical staining of tumor samples

Formalin-fixed paraffin-embedded human breast cancer tissues were obtained from US Biomax (BR1504, BR1505) and Shanghai Superchip Biotech (HBre-Duc052Bch, HBre-Duc068Bch-01, OD-CT-RpBre03-004). For xenografted tumors arising from MCF7 cells, tumors were extracted and formalin-fixed followed by paraffin-embedding. For immunostaining, paraffin sections were fixed in acetone, air dried, and then stored at −80°C. After abolishment of endogenous peroxidase activity (0.3% H2O2, 20 minutes), tissues were rehydrated with PBS and blocked with 3% BSA in PBS for 30 minutes. Then, sections were treated with primary antibodies specific for CD44 for 60 minutes at room temperature. Detection was achieved with compatible-conjugated secondary antibody (Invitrogen, CA, USA) and Horse radish-peroxidase-conjugated (HRP-conjugated) ABC amplification system.

### 4. CCK-8 assay

The proliferation of CD44^hi^ and Cd44^lo^ cells was determined by CCK-8 kit (Dojindo, Japan). Approximately purified 3.5×10^3^ cells in 100 ml were incubated in triplicate in 96-well plates. At 0, 24, 48, 72 and 96 h, the CCK-8 reagent (10 ml) was added to each well and incubated at 37°C for 2 h. The optical density at 450 nm was measured using an automatic microplate reader (Synergy4; BioTek, Winooski, VT, USA).

### 5. EdU assay

The cell proliferation was analyzed by EdU (5-ethynyl-20-deoxyuridine) assay using Cell-Light EdU DNA Cell Proliferation Kit (RiboBio, Shanghai, China). CD44^hi^ and CD44^lo^ cells (1×10^4^) were seeded in 96-well plates. After incubation for 48 h, 50 mM EdU was added and incubated at 37°C for another 2 h. Then, cells were fixed with 4% paraformaldehyde and stained with Apollo Dye Solution for proliferating cells. Finally, Hoechst 33342 was added to stain nucleic acids in all cells. The cell proliferation rate was calculated according to the manufacturer’s instructions. Images were taken using a fluorescence microscope (Nikon).

### 6. Mammosphere formation assay

Sphere formation assays were performed as described previously (Phillips et al, 2006). CD44^hi^ or CD44^lo^ cells from MCF7 or MMTV PyMT tumors were purified by FACS. To measure the mammosphere formation potential of the sorted cells, 1 × 10^3^ cells were plated in ultra-low-attachment 24-well dishes containing mammosphere growth medium (MEGM with all required growth factors as described above for the growth of HMLER cells plus B-27 supplements, without bovine pituitary extract (BPE]) from Lonza. After 12 days of culture at 37°C, the resulting mammospheres were counted.

### 7. In vivo tumor initiation assay

CD44^hi^ or CD44^lo^ cells from MMTV PyMT tumors were FACS sorted using Astrios System (Beckman). Purified cells were transplanted subcutaneously into NOD/SCID mice at concentrations of 100,000, 20,000, and 2000 cells per site to analysis of the tumor initiation potential. Cells were harvested in a suspension of 50% Matrigel (BD Biosciences, cat. no. 356231) in DME/F12 medium to a final volume of 50μl, then orthotropically injected into 4- to 6-week-old NOD/SCID mice. Tumor volume was measured at the time of sacrifice and calculated by the ellipsoid volume. 3-8 mice were used in each group. After 3-4 weeks the tumors were identified by palpation. Mice were killed after tumors reached a diameter of 1.5 cm, in accordance with institutional guidelines.

### 8. Live imaging of cell migration assay

Live imaging of cells was conducted using a Nikon Living-Cell Observer system with a confocal microscope (Nikon A1, Tokyo, Japan). In general, images were collected at 30-min or 1h intervals with exposure times of ~250 ms. Some of the movies were collected. Temperature was held at 37 °C and CO_2_ at 5%.

Migration experiments were performed according to the method described by Das et al.(Das et al, 2015). CD44^hi^ and CD44^lo^ cells from MCF7 cells were cultured on each dish separately with one silicone culture-insert (ibid) press-bonded to create confined wells (each well 3.25mm×7mm; growth area per well 22 mm^2^). When cells formed 100% confluent monolayers, the insert was removed to generate cluster migration in the forward facing direction with 0.5mm wound-gap. 3×10^4^ cells were seeded into each well with 80 μl cell culture medium and incubated at 37 °C in a 5% CO2 humidified incubator. The migration area of each cell= Cell area after 24 or 72 hours-Cell area at T0.

### 9. RNA Sequencing

Tumor tissue was enzymatically digested using mouse tumor dissociation kit (Miltenyi, Cat.130-096-730) before FACS analysis. Cells sorted from MMTV tumors with CD44^hi^/EpCAM^+^/CD45^-^ or CD44^lo^/EpCAM^+^/CD45^-^ features were collected. Total RNA was isolated using the Phenol-chloroform extraction and used for Illumina RNA sequencing analysis (Illumina HiSeq X-Ten). Transcriptomic libraries were constructed using Illumina TruSeq RNA Sample Preparation Kit (Illumina, San Diego, CA, United States). Reads with high-quality were aligned to the mouse reference mRNA (GRCh38) using Bowtie2. For gene ontology analysis, we used Gene Ontology (GO) and KEGG for gene functional and signaling pathway annotation. Hypergeometric test and the following FDR correction were used for enriched GO term or KEGG pathway analysis. After FDR corrected, significant enrichment was valued by the P-value cutoff of 0.05. The expression levels for each of the genes were normalized to fragments per kilobase of exon model per million mapped reads (FPKM) using RNA-seq by Expectation Maximization (RSEM). The hierarchical clustering was analyzed using Pearson correlation distance measure and average linkage method.

### 10. Quantitative real-time PCR

Real-time PCR was performed to validate gene expression in CD44^hi^ and CD44^lo^ cancer cells. Total RNA was extracted by RNeasy mini kit (Takara) according to the manufacturer’s instructions. Purified RNA was treated with RNase-free DNaseI to ensure complete degradation of potential genomic DNA contamination, followed by the addition of 25mM EDTA and heat denaturation of the enzyme. RNA samples were reverse transcribed to cDNA with random hexamer primers using a SuperScript III First-Strand cDNA Synthesis Kit (Takara). Real-time PCR primers were designed with RealTime Design software (http://www.bio-searchtech.com) and were custom made by Invitrogen.

### 10. Analysis of CD44 expression

Cells were harvested and washed with Hanks' balanced salt solution (HBSS) supplemented with 2% bovine serum albumin, (pH 7.4). A single cell suspension (10^6^ cells/ml) was incubated with CD44-PE antibody or isotope control antibodies on ice for 1 h. Cells were washed three times and analyzed using a flow cytometer (Beckman-Coulter, Brea, USA), and at least 8,000 cells were analyzed per sample. All experiments were performed at three times.

### 11. Analysis of CD44 expression pattern

Cells were harvested and total RNA was purified using TRIzol reagent. For reverse transcription, a random nonamer-oligo dT combination was used as the primer. To determine the CD44 expression pattern, we use a human-specific primer pair spanning the entire variant region (forward primer: 5’-AGT CAC AGA CCT GCC CAA TGC CTT T, reverse primer: 5’-TTTGCT CCA CCT TCT TGA CTC CCA TG-3’(Banky et al, 2012; Yang et al, 2015). All CD44 variants are theoretically amplified. The PCR reaction mixture was obtained from Takara. The PCR cycling procedures were described as our previous report(Yang et al, 2015).

### 12. The shift of CD44 in vivo

To determine the generation of CD44^hi^ cells from CD44^lo^ cells and vice versa in vivo, 1×10^6^ CD44^hi^, CD44^lo^, or the mixer cells (CD44^hi^:CD44^lo^=1:1) obtained from MMTV PyMT tumors, were transplanted orthotropically into NOD/SCID mice. The tumors were removed 4 weeks after transplantation, and digested into single-cell suspensions using mouse tumor dissociation kit (Miltenyi, Cat.130-096-730). The cell suspension was analyzed for CD44 expression using an Astrios flow cytometer.

### Statistical analysis

Statistical analysis was performed using GraphPad InStat software (GraphPad Software, Inc.). Nonparametric Mann-Whitney tests were performed to measure the differences in CD44 expression, tumor formation, mammosphere-forming capacity, and invasiveness between CD44^hi^ and CD44^lo^ cells. The significance of differences among groups was determined by one-way ANOVA t test, or Fisher exact test accordingly. Statistically significance was considered as P < 0.05.

## Author Contributions

Conceptualization, CX.Y, ML. C., F.G.; Formal Analysis, T.M., R. J., J.V.; Investigation, YW.L., YQ.H., Y.D., GL. Zh.; Resources, CX.Y., YW.L., YQ.H., YD., GL. Zh., F.G; Funding Acquisition, CX.Y., F.G.

## Acknowledgments

This work was supported by the National Natural Science Foundation of China (81672843, 81572821, 81502490, 81502491, 81402419), Shanghai Municipal Education Commission-Gaofeng Clinical Medicine Grant Support (20171924), the Natural Science Foundation of Shanghai Municipality (14YF1412200), and the Shanghai Shen-Kang Hospital Development Center (SHDC22014004).

## Supplemental Figures Legends

**Fig. S1.**
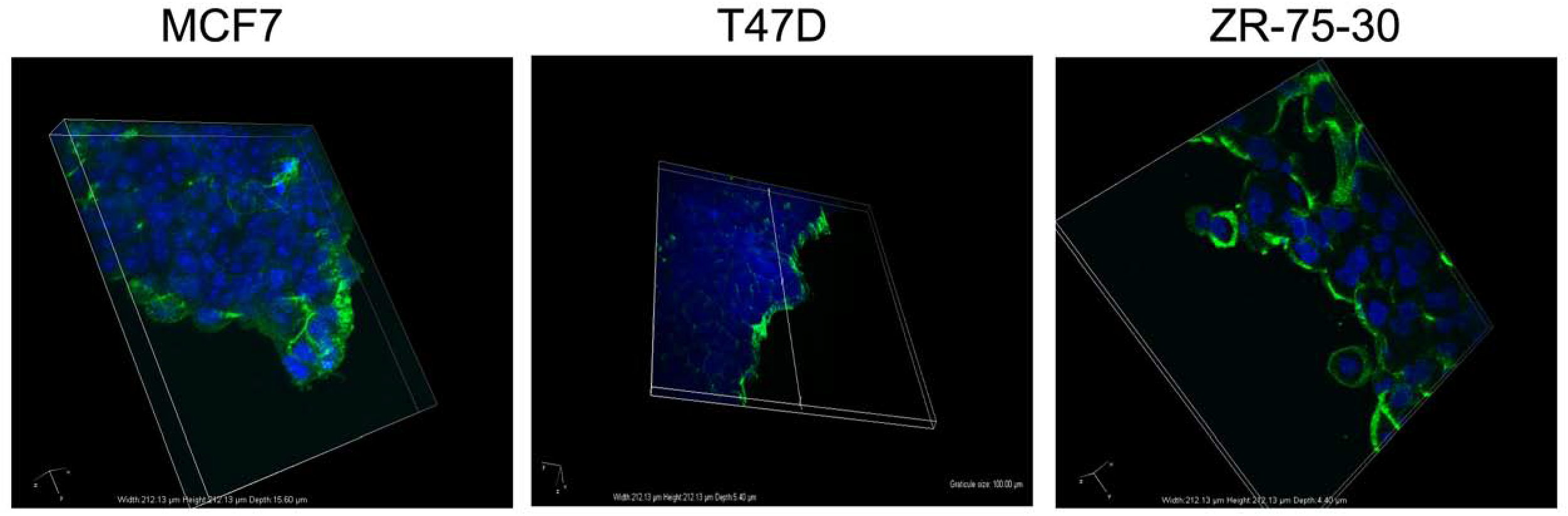
Collective migration of breast cancer cells in 2-D monolayer culture. Confocal XYZ vertical images showed that CD44 maintained in the leading edge of collective cell clusters.

**Fig. S2.**
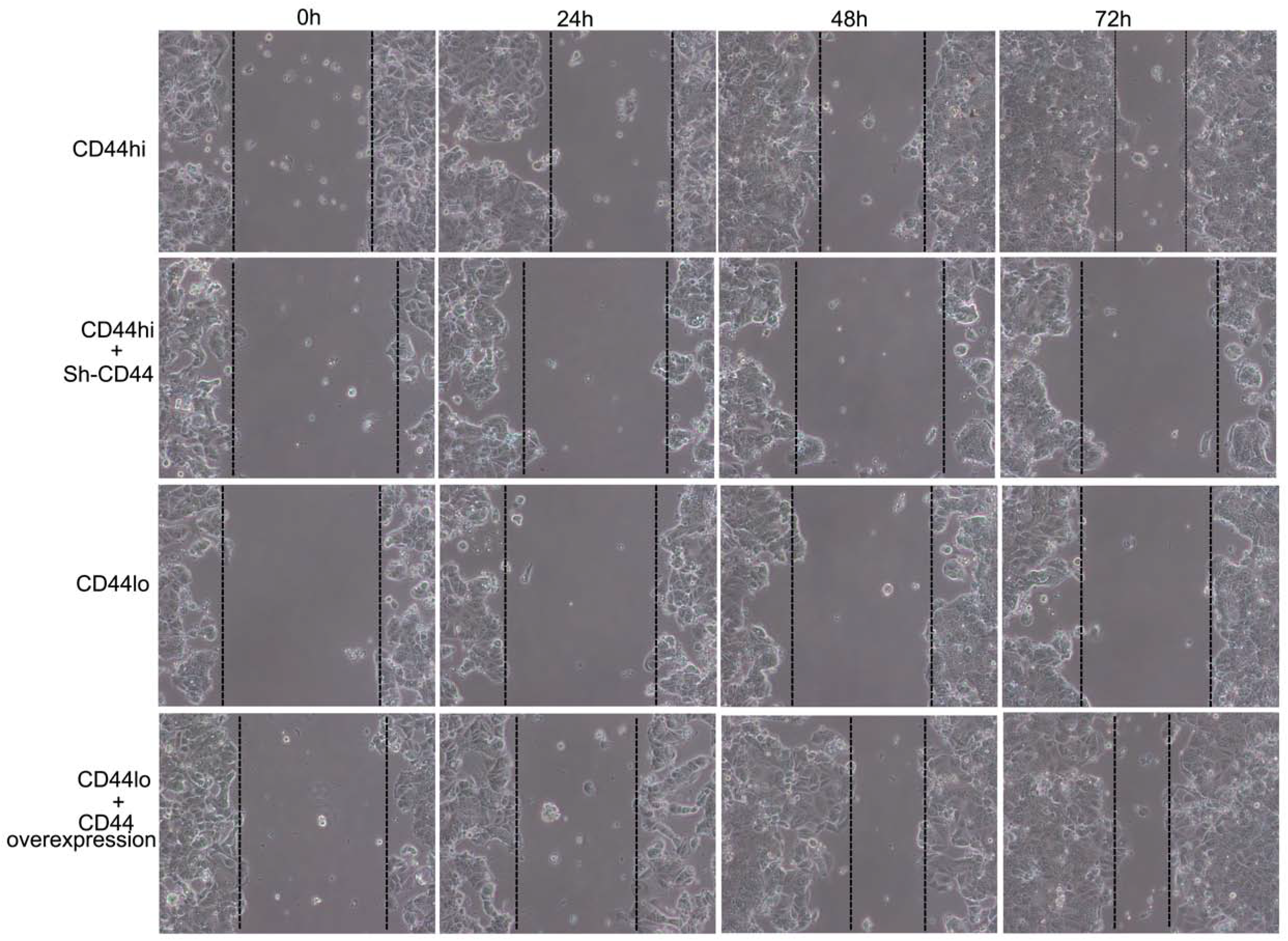
The role of CD44 in mediating collective migration in monolayer sheets. Purified CD44hi or CD44lo cells obtained from MCF7 cells were transfected with Sh-CD44 virus or CD44 virus, separately, then examined for the wound closure. Cells were cultured in 6-well plates until they formed 100% confluent monolayers. Then a 100 mm “wound” scratches were made using a sterile pipette tip. The cells was incubated with MEM containing 1% FBS for different periods. Then the pictures were captured.

**Fig. S3.**
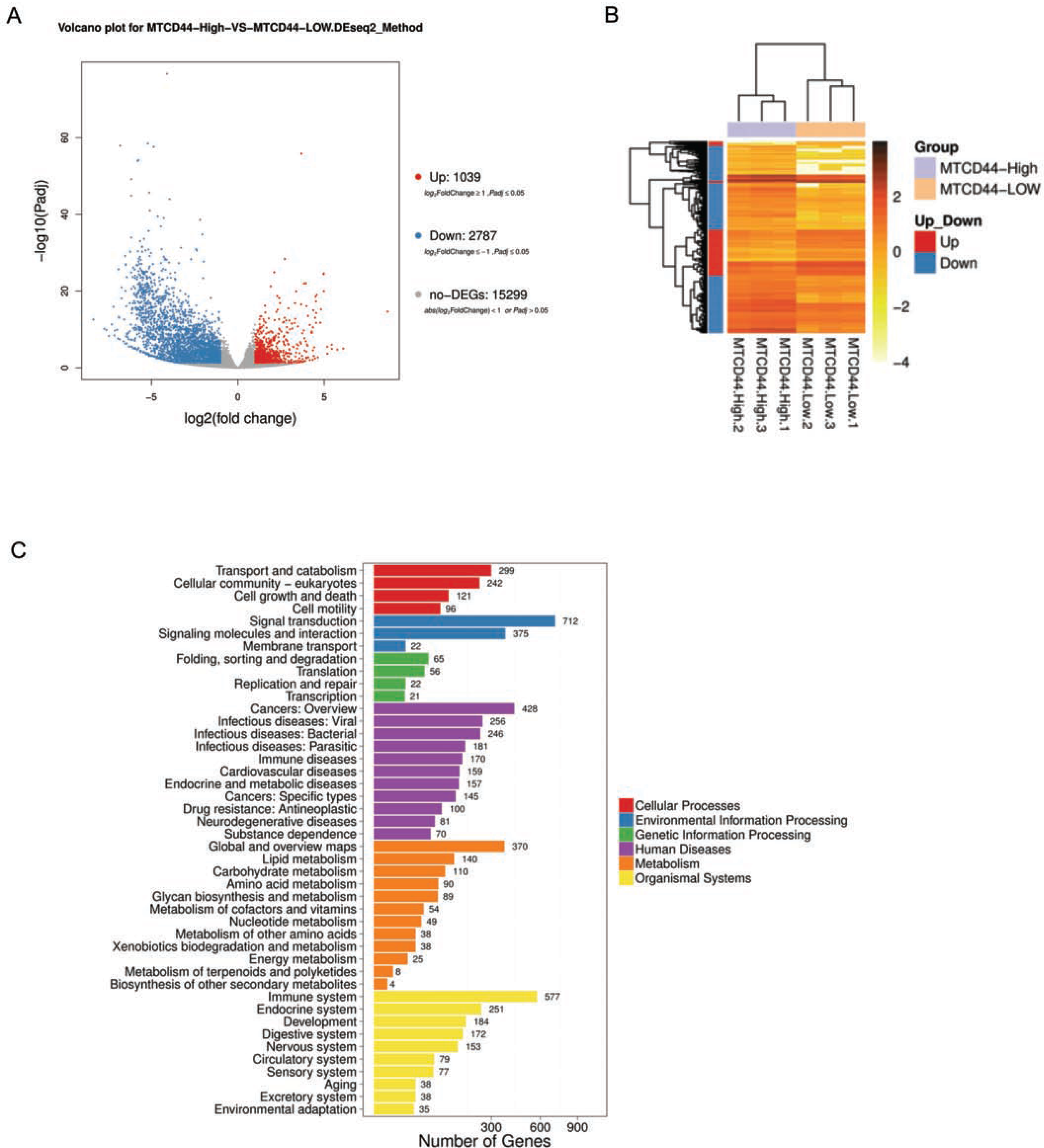
Gene expression of purified CD44hi/leading and CD44lo/following cells. (A) Scatter plot showing, expression differences greater in follower cells(CD44lo) (y-axis) versus leader cells (CD44hi) (x-axis). (B) Unsupervised hierarchical clustering of 1422 significantly differentially expressed genes in leader (purple, N=2782) and follower (yellow,N=1039) cells. The hierarchical clustering was performed using Pearson correlation distance measure and average linkage method. The log2 transformed gene levels are represented after quantile normalization. (C)Bioinformatic analysis of gene expression differences in CD44hi/leading and CD44lo/following cells.

**Fig. S4.**
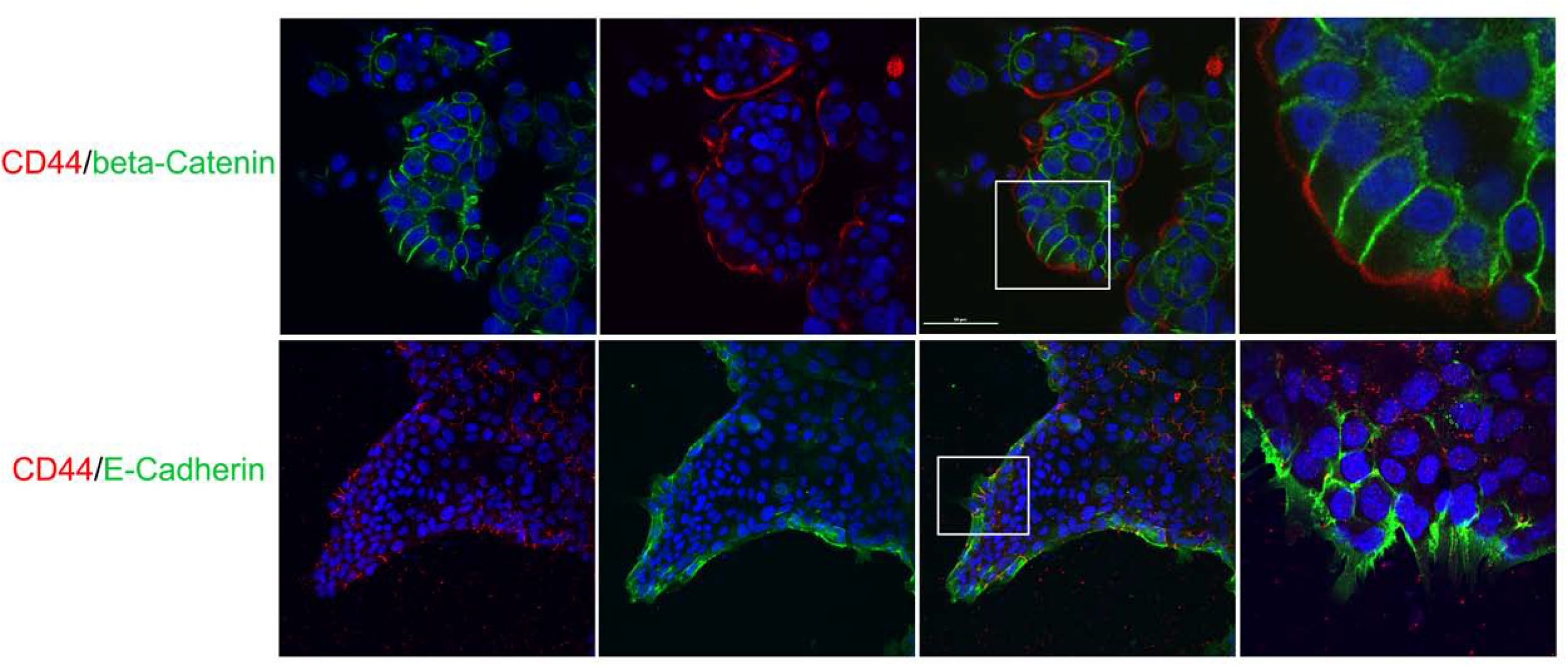
Confocal images of co-localization of CD44 and E-cadherin, or β-catenin, during collective migration of MCF7 cells. Co-localization of CD44 and E-cadherin, or β-catenin, in migrating monolayer. MCF7 cells or EpCAM^+^/CD45^-^ primary cancer cells purified from MMTV/PyMT mice, were cultured in the wound chamber (ibid). A sheet migration of epithelial cells was generated as described before. Then, cells were fixed for CD44 and E-cadherin, or β-catenin expression at 24h post wounding by immunofluorescence assay. Representative images were presented. Scale bar, 50 μm.

**Fig. S5.**
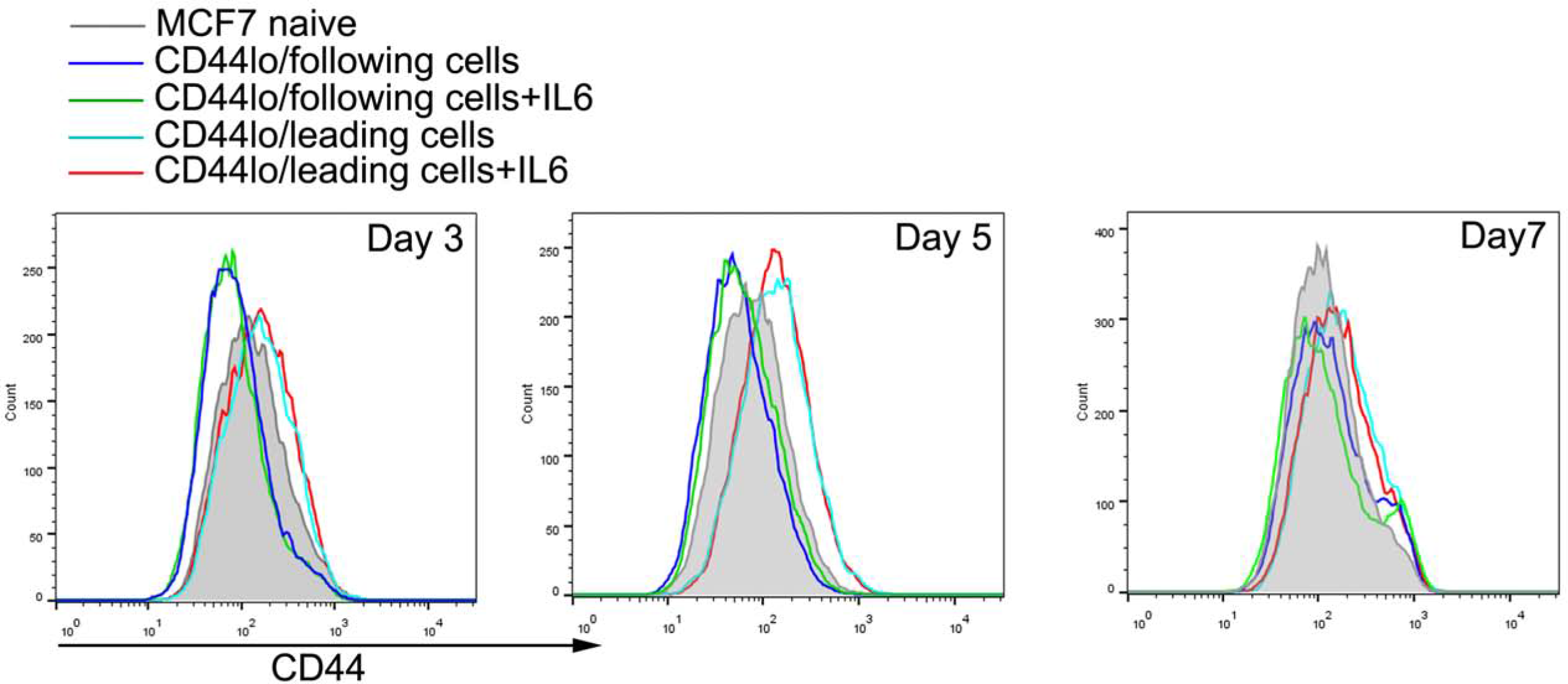
Flow cytometric analysis of changes of CD44 levels after IL6 stimulation. Flow cytometric analysis of changes of CD44 levels after stimulated with rhIL-6. Purified CD44^hi^ or CD44^lo^ cells obtained from MCF7 cells were incubated in the absence or presence of rhIL-6 (50ng/ml) for 0-7days, then examined for CD44 levels by FACS.

**Fig. S6.**
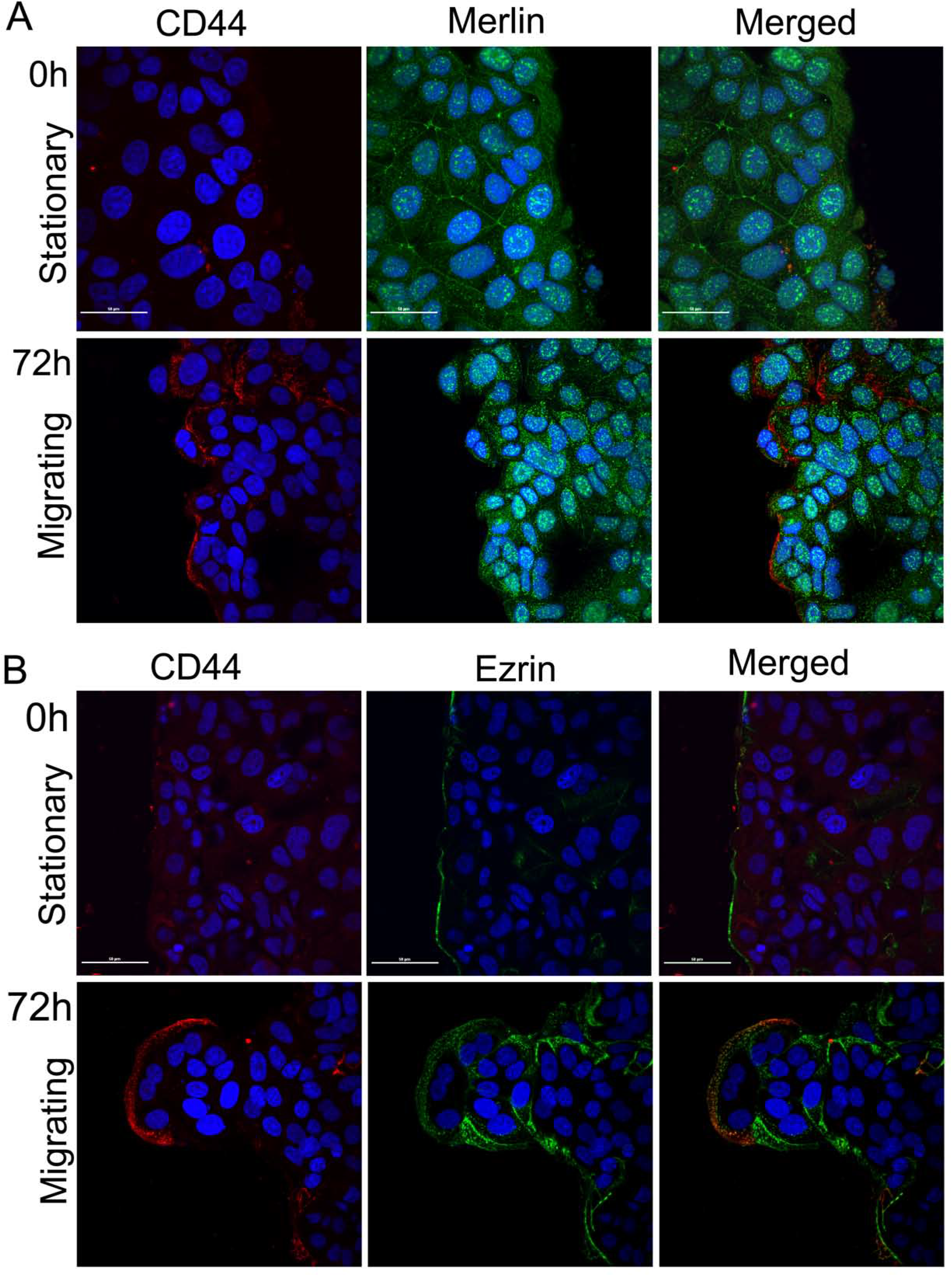
Co-localization of CD44 and Merlin or Ezrin in stationary and migrating cells during collective migration of MCF7 cells. (A) Co-localization of CD44 and Merlin in stationary (upper) and moving (Lower) monolayer. CD44^hi^ or CD44^lo^ MCF7 cells were cultured in the wound chamber (ibid). A sheet migration of epithelial cells was generated and fixed for Merlin and CD44 expression at 0h and 72h post wounding by immunofluorescence assay. Representative images were presented. Scale bars, 50 μm. (B) Localization of CD44 and ezrin in stationary (upper) and migrating ((Lower) monolayer, ezrin and CD44 expression at 0h and 72h post wounding were determined by immunofluorescence assay. Scale bars, 50μm.

**Table.S1.** Correlation between CD44 expression and the clinico-pathological characteristics in the studied cohort

| Tissue marker | n | CD44 expression (IOD) |  | Significance |
| --- | --- | --- | --- | --- |
|  |  | Median | 95% CI | *P* value |
| Age |  |  |  |  |
| <50 y | 97 | 4927 | 3892-5962 | 0.2802 |
| ≥50 y | 150 | 6599 | 5558-7640 |  |
| Tumor size |  |  |  |  |
| <2 cm | 108 | 4775 | 3771-5779 | 0.4856 |
| ≥2 cm | 149 | 5814 | 4767-6861 |  |
| Histologic grade |  |  |  |  |
| 1 | 53 | 2522 | 1070- 10256 | 0.4792 |
| 2 | 134 | 2755 | 4126- 7176 |  |
| 3 | 60 | 2537 | 2984- 13207 |  |
| Lymphnode metastasis |  |  |  |  |
| Yes | 54 | 2522 | 2819- 7388 | 0.5405 |
| No | 193 | 2680 | 4511- 7854 |  |
| ER |  |  |  |  |
| Negative | 112 | 3610 | 6290-11240 | <0.0001* |
| Positive | 135 | 1930 | 2206-3017 |  |
| PgR |  |  |  |  |
| Negative | 147 | 3588 | 6032-10621 | <0.0001* |
| Positive | 100 | 1890 | 2120-2986 |  |
| HER2 |  |  |  |  |
| Negative | 126 | 3318 | 5744-11282 | 0.0005* |
| Positive | 135 | 1999 | 2918-4234 |  |
| Molecular subtype |  |  |  | < 0.0001* |
| Luminal-like A | 41 | 2060 | 1754- 3154 |  |
| Luminal-like B | 92 | 1963 | 2249- 3186 |  |
| HER2 over-expressing | 44 | 2790 | 3787- 7175 |  |
| Triple negative | 70 | 4866 | 7677- 17077 |  |

\*Indicated statistical significance (p <0.05)

## Supplemental Videos Legends

Movie 1: The migration rate of CD44^hi^ cells and CD44^lo^ cells from 0h to 17h

Movie 2: The migration rate of CD44^hi^ cells and CD44^lo^ cells from 42h to 72h

Movie 3: CD44^hi/DsRed-^ subset leads the migration of CD44^lo/DsRed+^ subset purified from MCF7 cells from 24h to 38h

Movie 4: CD44^hi/DsRed-^ subset leads the migration of CD44^lo/DsRed+^ subset purified from MCF7 cells from 39h to 55h

Movie 5: CD44^hi/DsRed+^ subset leads the migration of CD44^lo/DsRed-^ subset purified from T47D cells from 48h to 72h

Movie 6:The CD44^lo^/following cells sorted from MCF7 cells, carrying a CD44 promoter biosensor in which YFP reports on CD44 gene expression, can spontaneously switch to CD44^hi^ state according to collective migration.

Movie 7:The CD44^hi^/following cells sorted from MCF7 cells, carrying a CD44 promoter biosensor in which YFP reports on CD44 gene expression, can spontaneously switch to CD44^lo^ state according to collective migration.

## Notes

Disclosure of Potential Conflicts of Interest: No potential conflicts of interest were disclosed.

